# Genomic landscape of drug response reveals novel mediators of anthelmintic resistance

**DOI:** 10.1101/2021.11.12.465712

**Authors:** Stephen R. Doyle, Roz Laing, David Bartley, Alison Morrison, Nancy Holroyd, Kirsty Maitland, Alistair Antonopoulos, Umer Chaudhry, Ilona Flis, Sue Howell, Jennifer McIntyre, John S. Gilleard, Andy Tait, Barbara Mable, Ray Kaplan, Neil Sargison, Collette Britton, Matthew Berriman, Eileen Devaney, James A. Cotton

**Affiliations:** Wellcome Sanger Institute; Hinxton, United Kingdom; Institute of Biodiversity Animal Health and Comparative Medicine, College of Medical, Veterinary and Life Sciences, University of Glasgow; Glasgow, United Kingdom; Moredun Research Institute; Penicuik, United Kingdom; Royal (Dick) School of Veterinary Studies, University of Edinburgh; Edinburgh, United Kingdom; Department of Infectious Diseases, College of Veterinary Medicine, University of Georgia; Athens, United States; Department of Comparative Biology and Experimental Medicine, Host-Parasite Interactions Program, Faculty of Veterinary Medicine, University of Calgary; Calgary, Canada

## Abstract

Understanding the genetic basis of anthelmintic drug resistance in parasitic nematodes is key to improving the efficacy and sustainability of parasite control. Here, we use a genetic cross in a natural host-parasite system to simultaneously map resistance loci for the three major classes of anthelmintics. This approach identifies novel alleles for resistance to benzimidazoles and levamisole and implicates the transcription factor, *cky-1*, in ivermectin resistance. This gene is within a locus under selection in ivermectin resistant populations worldwide; functional validation using knockout experiments supports a role for *cky-1* overexpression in ivermectin resistance. Our work demonstrates the feasibility of high-resolution forward genetics in a parasitic nematode, and identifies variants for the development of molecular diagnostics to combat drug resistance in the field.

**One-Sentence Summary:** Genetic mapping of known and novel anthelmintic resistance-associated alleles in a multi-drug resistant parasitic nematode

## Main Text

Over a billion people and countless livestock and companion animals require at least annual treatment with anthelmintic drugs to control parasitic worm (helminth) infections. The rapid and widespread evolution of resistance to these drugs is a significant health concern in livestock (*1*) and places an economic burden on food production. Resistance is present on every continent where anthelmintics are used; in many places, individual drug classes are now ineffective, and some farms have resistance to every major class of drug (*2*), threatening the economic viability of livestock farming. In Europe, gastrointestinal helminths of livestock are responsible for annual production losses of €686 million, of which €38 million is associated with anthelmintic resistance (*3*). Drug resistance is also now a major concern in the treatment of helminths infecting dogs (*4, 5*), with multiple drug resistance to all major anthelmintic classes in the dog hookworm now common in the USA (*6*). The same classes of drugs to which veterinary parasites have rapidly evolved resistance are also used to control related human-infective helminths, which are targeted at scale by some of the largest preventative chemotherapy programmes in the world. Although less established in human-infective helminths, the emergence of widespread anthelmintic resistance – echoing the current global emergency around antimicrobial resistance – will have serious socio-economic and welfare impacts on people infected with parasitic worms and derail hard-won progress towards the proposed eradication and elimination of helminths over the next decade (*7, 8*).

Despite extensive efforts, the causal mutations and mechanisms of resistance in parasitic helminths remain largely unresolved. Many candidate “resistance genes” have been proposed for most drug classes; these candidates are primarily homologues of genes that confer resistance in the free-living model nematode *Caenorhabditis elegans,* and are subsequently assayed for differences in genetic variation and/or gene expression in parasite isolates that vary in their response to treatment (*9–11*). A successful example of this approach is the identification of variants of β-tubulin that inhibit tubulin-depolymerisation by benzimidazole-class anthelmintics (*12, 13*). These variants, particularly at amino acid positions 167, 198 and 200 of β-tubulin isotype 1 (*14–16*), have subsequently been shown to be associated with resistance in many parasitic species for which benzimidazoles have been extensively used, and a number of these parasite-specific mutations have been functionally validated in *C. elegans* (*17, 18*). However, these three variants are unlikely to explain all phenotypic variation associated with resistance (*19, 20*), and it is unknown to what degree other variants contribute to benzimidazole resistance in parasitic species. For other drug classes, few candidate genes have been functionally validated or shown to be important in natural parasite populations. For example, concurrent mutation of three glutamate-gated chloride channels (*glc-1, avr-14, avr-15*) enables resistance to high levels of ivermectin by *C. elegans* (*21*), yet no strong evidence of selection on these channels in any parasitic species has been demonstrated to date. On the one hand, the many genes proposed may reflect that resistance is a complex, quantitative trait where similar resistance phenotypes can be derived from variation in multiple loci. Alternatively, resistance may be species and/or population-specific, and evolve independently under subtly different selection pressures (*22*). However, some candidates are likely to have been falsely associated with resistance, as most studies present relatively weak genetic evidence from the analysis of single or few candidate loci in small numbers of helminth populations that often differ in both drug susceptibility and geographic origin. Many helminth species are exceptionally genetically diverse (*23–26*), and consequently, candidate gene approaches have limited power to disentangle causal variation from linked but unrelated background genetic variation, a situation that is exacerbated by the experimental intractability and inadequate genomic resources available for many parasitic helminths (*9*).

Here we describe a genome-wide forward genetics approach using the parasitic nematode *Haemonchus contortus* as a model to identify genetic variation associated with resistance to three of the most important broad-spectrum anthelmintic drugs globally: ivermectin, levamisole, and benzimidazole. *H. contortus* is an economically important gastrointestinal parasite of livestock and one of only a few genetically tractable parasites used for drug discovery (*27, 28*), vaccine development (*29, 30*), and anthelmintic resistance research (*22*). Our approach has exploited a genetic cross between the susceptible MHco3(ISE) and multi-drug resistant MHco18(UGA) strains of *H. contortus* (**fig. S1 A**), allowing us to investigate resistance in a natural host-parasite system while controlling for confounding genetic diversity that differentiates parasite strains (see **Supplementary materials** regarding the establishment and validation of the cross). Using an eXtreme Quantitative Trait Locus (X-QTL) (*31, 32*) analysis framework, whereby pools of F3-generation progeny from F2 adults treated *in vivo* were sampled pre- and post-treatment for each drug (**fig. S1 B**; n = 3 parasite populations per drug class maintained in independent sheep; **fig. S2**) and analysed by whole-genome sequencing (**table S1**), we aimed to identify drug-specific quantitative trait loci (QTLs) associated with resistance throughout the genome. These QTLs and specific variants were independently validated using genome-wide variation from populations of *H. contortus* obtained from ten US farms of known resistance phenotype (see **Supplementary materials** for a description of the US farms and quantitative phenotyping; **table S2, fig. S3, fig. S4**), and from more than 350 individual parasites sampled throughout the world where *H. contortus* is endemic (*25, 33*).

### A genetic cross between genetically-distinct susceptible and multi-drug resistant strains reveals drug-specific QTL after selection

A key feature and thus advantage of using a genetic cross to map anthelmintic resistance loci is that the high degree of within-strain diversity and genome-wide genetic divergence is controlled by admixture in the F1 generation of the cross. The susceptible and resistant parental *H. contortus* strains of the cross are highly genetically differentiated throughout their genomes (**Fig. 1A**; mean *F*_ST_ = 0.089 ± 0.066 SD; n = 16,794,366 single nucleotide variant sites), typical of two parasite strains sampled from different continents (*25, 34*). In subsequent generations, both susceptible and resistant alleles segregate at moderate frequencies in the absence of selection, and genetic recombination breaks down the linked genetic variation that defines and differentiates the parental strains. This was evident by a significantly lower genome-wide genetic differentiation in the F3-generation control population (genome-wide mean *F*_ST_ = 0.012 ± 0.004) and absence of discrete peaks of high genetic differentiation (**Fig. 1B: Control**). In contrast, after each drug treatment, discrete QTLs that differ between each drug class were revealed: after benzimidazole treatment, we identified a major QTL on chromosome 1 (**Fig. 1B: Benzimidazole**); after levamisole, two QTLs on chromosome 4 and 5 (**Fig. 1B: Levamisole**); and after ivermectin, a major QTL on chromosome 5 and minor QTLs on chromosomes 2 and 5 (**Fig. 1B: Ivermectin**).

**Fig. 1.**
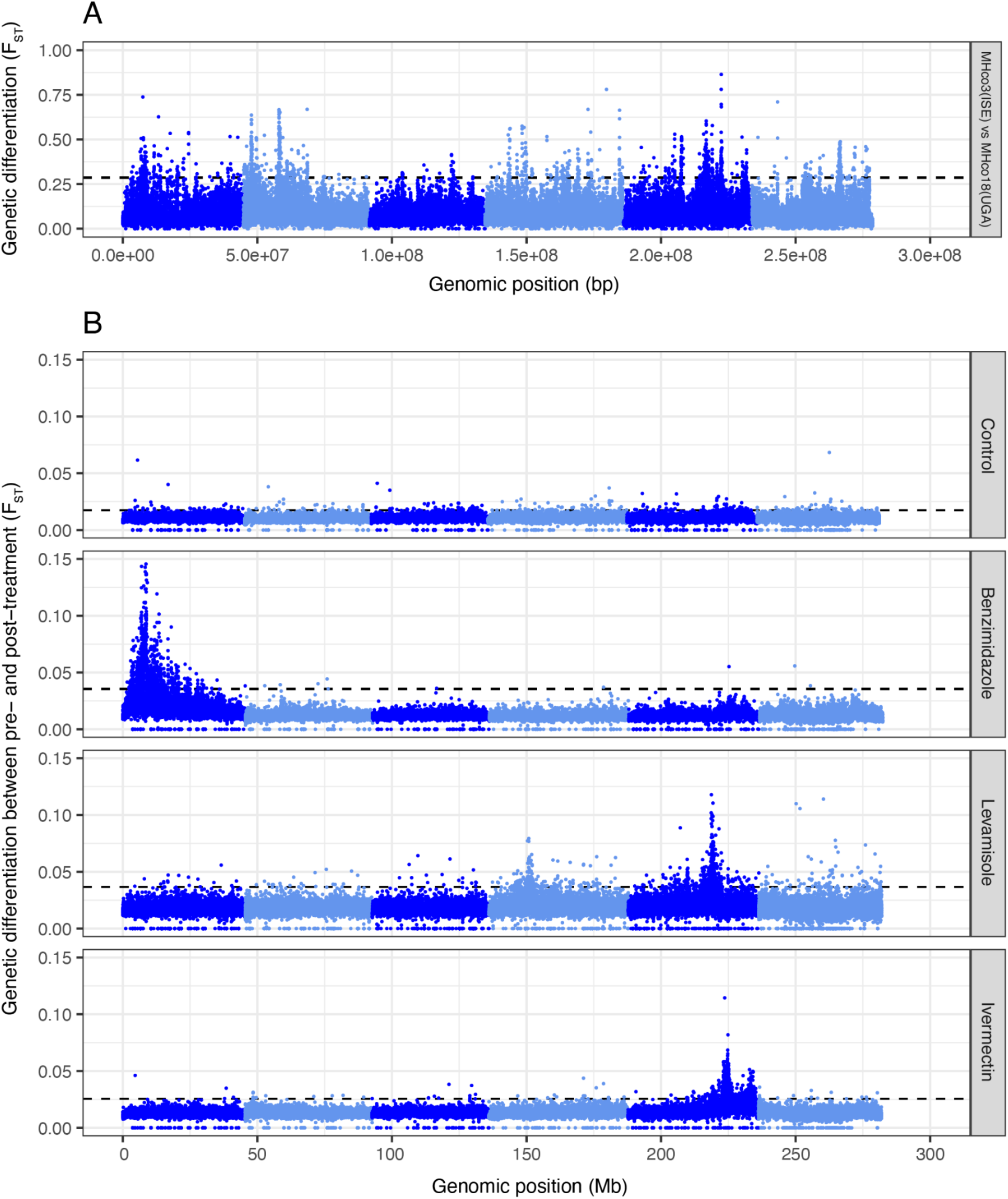
A genetic cross followed by drug selection reveals discrete QTLs associated with each anthelmintic drug class. (**A**) Genome-wide comparison of susceptible MHco3(ISE) and multidrug-resistant MHco18(UGA) parental strains revealed broad-scale genetic differentiation on all chromosomes. (**B**) Comparison of genome-wide differentiation between F3 generation pooled infective-stage larvae (L3, n = 200) sampled pre- and post-treatment revealed distinct genomic regions or QTLs associated with benzimidazole, levamisole, and ivermectin drug treatment. An untreated control where sampling was time-matched to the treated groups is shown for comparison. In all plots, each point represents the mean genetic differentiation (*F*ST) from three biological replicates in five kb sliding windows, and the dashed line represents the genome- wide mean *F*ST + 3 SD for each comparison (See **fig. S2** for genome-wide replicate data). Individual chromosomes are indicated by alternating dark and light blue shading.

### Variation at β-tubulin isotype 1 and a novel β-tubulin isotype 2 variant is associated with high levels of benzimidazole resistance

The β-tubulin isotype 1 (HCON_00005260) gene and, in particular, nonsynonymous changes at coding positions 167, 198, and 200 have been widely associated with benzimidazole resistance in *H. contortus* (*13, 15, 16, 35*) and other nematodes frequently exposed to benzimidazole treatment (*17, 36*). After benzimidazole selection, a single broad QTL was found on chromosome 1 (**Fig. 2A**; see **Supplementary materials** for further discussion of the genetic structure of the QTL) containing the β-tubulin isotype 1 locus. Within this gene, we identified a significant increase in the frequency of a Phe200Tyr variant (a phenylalanine [reference susceptible variant] to tyrosine [resistant variant] substitution at position 200) from pre- to post-treatment and relative to untreated controls (**Fig. 2B**; *P* = 1.7e-26, genome-wide Cochran–Mantel–Haenszel (CMH) test between replicates). We also identified a small increase in frequency of the Phe167Tyr variant (mean *freq*_pre-treatment_ = 0.14 to *freq*_post-treatment_ = 0.20), however, no variation was found at the Glu198 position. Considering the previous association between these variants and benzimidazole resistance, we conclude that the Phe200Tyr variant is the primary driver of phenotypic resistance in the X-QTL population.

**Fig. 2.**
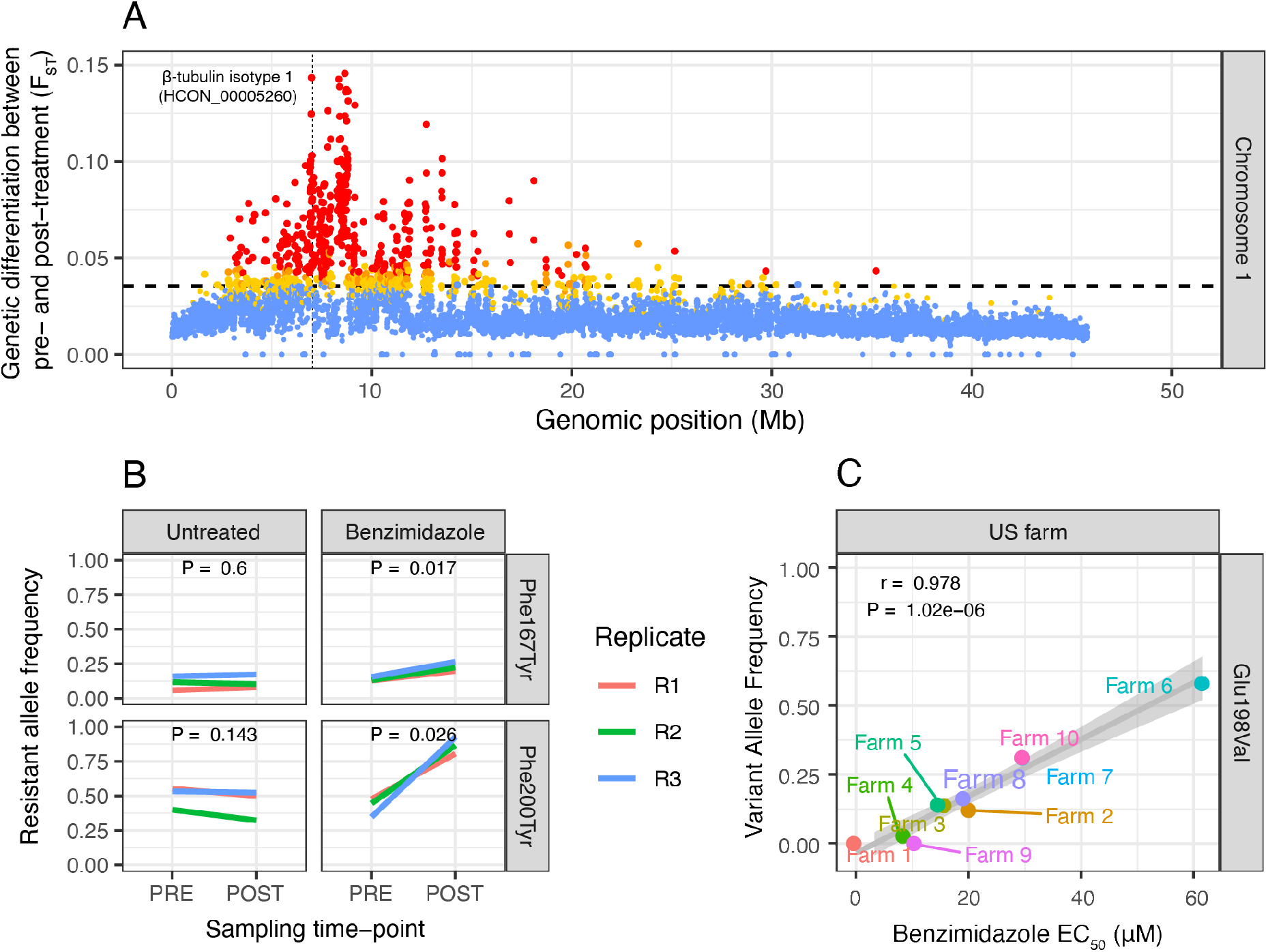
Characterisation of QTL associated with benzimidazole resistance. **(A)** Chromosome-wide genetic differentiation between pre- and post-benzimidazole treatment on chromosome 1. Each point represents the mean *F*ST in a five kb window; points are coloured based on the concordance of individual replicates indicated by none (blue), 1 of 3 (yellow), 2 of 3 (orange), or all 3 (red) above the genome-wide threshold. The genome-wide threshold is defined as the mean + 3 SD of the chromosome-wide *F*ST indicated by the horizontal dashed line, whereas the vertical dashed line highlights the position of the β-tubulin isotype 1 (HCON_00005260) gene. **(B)** Allele frequency change at Phe167Tyr and Phe200Tyr variant positions of β-tubulin isotype 1 pre- and post-treatment, including untreated time-matched control. Coloured lines show biological replicates. *P*-values are calculated using pairwise t-tests of allele frequency by sampling time point (i.e., pre- and post-treatment). **(C)** Correlation between benzimidazole EC50 concentration (μM) observed at particular farms and Glu198Val variant frequency of β-tubulin isotype 2 (HCON_00043670) on US farms. Pearson’s correlation (r) and associated *P*-value together with the trendline and standard error of the linear regression are shown.

*Haemonchus contortus* has multiple β-tubulin genes (*37*), and deletion of the β-tubulin isotype 2 gene (HCON_00043670) on chromosome 2 has been associated with increased levels of resistance beyond that of mutations in the isotype 1 gene alone (*14*). Here, we found no evidence of deletions in isotype 2. However, a minor but not significant increase in genetic differentiation between pre- and post-treatment populations was found at this locus, and a Glu198Val variant at a homologous site to a known resistance variant in isotype 1 was present at a low frequency in the genetic cross (*freq*_pre-treatment_ = 0.260 to *freq*_post-treatment_ = 0.323; not significant genome-wide CMH). However, on the US farms, the Glu198Val variant did vary in frequency between farms and was significantly correlated (r = 0.978, *P* = 1.02e-6; Pearson’s correlation) with EC_50_ values for benzimidazole resistance (**Fig. 2C**). The variance observed in EC_50_ among resistant farm populations was not caused by variation in the frequency of the Phe200Tyr mutation of the isotype 1 gene, as this variant was already at high frequency in all populations, except for the farm that was susceptible to benzimidazoles (Farm 1; **fig. S5**). These data suggest that once the isotype 1 Phe200Tyr variant has reached near fixation in the population, the Glu198Val variant of isotype 2 mediates higher levels of benzimidazole resistance than conferred by the Phe200Tyr variant alone. As such, this novel allele present in β-tubulin isotype 2 should be considered, in addition to the well-characterised isotype 1 variants, as a genetic marker for benzimidazole resistance.

In addition to the association with benzimidazole resistance, it has been suggested that the β-tubulin isotype 1 Phe200Tyr variant in *H. contortus* (*38–40*) and also at an equivalent variant site in a β-tubulin gene in the human-infective filarial nematode *Onchocerca volvulus* (*41*) is associated with ivermectin resistance. Here we found no evidence of selection on either the Phe167Tyr or Phe200Tyr variants (or any variant found in the region) in X-QTL analyses of ivermectin treatment (**fig. S6A**), nor any correlation with ivermectin EC_50_ on the US farms (**fig. S6B**). These data reaffirm that mutations in β-tubulin isotype 1 are specific to benzimidazole resistance.

### Levamisole selection implicates acetylcholine receptors, including a novel acr-8 variant, with resistance

The anthelmintic activity of levamisole is due to its antagonistic effect on nematode nicotinic acetylcholine receptors (*42*), and resistance in *C. elegans* is typically associated with variation in subunits of these receptors or other accessory proteins that contribute to acetylcholine-mediated signalling (*43*). Here we identified two major QTLs on chromosomes 4 and 5 that contain a tandem duplication of the acetylcholine receptor subunit β-type *lev-1* (HCON_00107690 & HCON_00107700) and acetylcholine receptor subunit *acr-8* (HCON_00151270), respectively (**Fig. 3A**).

**Fig. 3.**
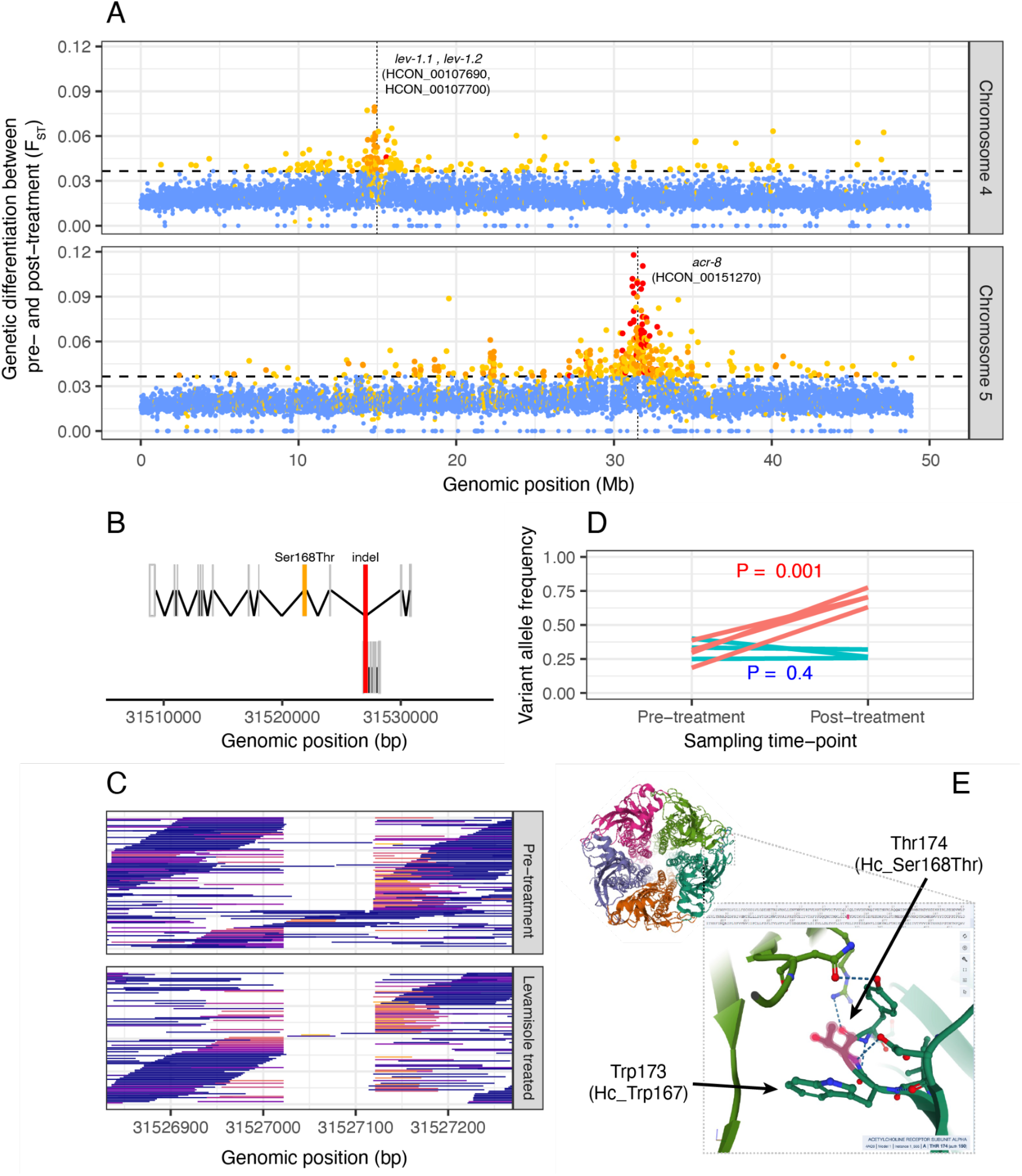
Characterisation of QTL associated with levamisole resistance. (**A**) QTL between pre-treatment and levamisole-treated parasites on chromosome 4 (top) and chromosome 5 (bottom). Each point represents the mean *F*ST in a five kb window; points are coloured based on the concordance of individual replicates indicated by none (blue), 1 of 3 (yellow), 2 of 3 (orange), or all 3 (red) above the genome-wide threshold (horizontal dashed line; mean + 3 SD of the chromosome-wide *F*ST). (**B**) Gene model for *acr-8* (top; HCON_00151270) and a cuticle collagen (bottom; HCON_00151260), highlighting the position of the overlapping *acr-8*/levamisole-associated indel and the Ser168Thr variant of *acr-8*. (**C**) Visualisation of sequencing reads supporting the *acr-8* intronic indel. Mapped reads are coloured to reflect the degree to which they have been clipped to allow correct mapping in the presence of the deletion, i.e. reads that have not been clipped are blue, whereas reads that are moderate to highly clipped are coloured red to yellow, respectively. (**D**) Comparison of Ser168Thr variant frequency between pre- and post-levamisole treatment (red) and time-matched untreated controls (green). (**E**) Structure of the pentameric cys-loop acetylcholine receptor of *Torpedo marmorata* (Protein Data Base ID: 4), one of the few species from which the receptor’s structure has been resolved (*55*). The Trp[Ser/Thr]Tyr motif is highly conserved among the clade V nematodes (**fig. S7**) and the distantly related alpha subunit of *T. marmorata*; Thr174, the homologous position of the *H. contortus* Hc_Ser168Thr variant of *acr-8,* lies within the acetylcholine binding pocket at the interface of the alpha and gamma subunits and adjacent to Trp173 (*H. contortus* Hc_Trp167), a residue essential for ligand binding.

The *H. contortus acr-8* gene (**Fig. 3B**) has long been implicated in levamisole resistance; a truncated isoform of *acr-8* containing the two first exons and a part of intron 2 (previously called *Hco-acr-8b*) (*44*), and subsequently, a 63 bp indel between exons 2 and 3 have been associated with resistance based on their presence in several resistant isolates (*45*). However, the functional consequence of these variants in mediating levamisole resistance *in vivo* is not yet clear. Here, we identified two larger deletion variants spanning 31,527,022 to 31,527,119 (97 bp) or 31,527,121 (99 bp) on chromosome 5 that increased in frequency from 73.47% in the pre-treatment population to 86.58% after levamisole treatment (**Fig. 3C**; paired t-test across replicates, *P* = 0.1). However, the *acr-8* indel was present in the levamisole susceptible parental MHco3(ISE) strain (59.05%) and was present at only a slightly higher frequency in the resistant MHco18(UGA) strain (63.55%). Thus, these data argue that the *acr-8* indel is a poor marker of levamisole resistance.

We did, however, identify a nonsynonymous variant (Ser168Thr) in *acr-8* that was strongly correlated with resistance across multiple datasets. In the X-QTL analyses, Ser168Thr increased to a high frequency after drug selection in the F2 generation (**Fig. 3D**; position 31,521,884; genome-wide CMH: *P* = 1.6e-15; allele frequency change pre- vs post-treatment: *P* = 1.0e-4; in time-matched no-treatment control: *P* = 0.4). It was also found at a high frequency in the USA field population with the highest levamisole drug resistance phenotype (Farm 7; *freq*_Ser168Thr_ = 0.64). This association was supported in global diversity data of *H. contortus* (*25*), where we found the Ser168Thr variant fixed in parasites from the Kokstad (KOK; South Africa) population (*freq*_Ser168Thr_ = 1.0; n = 4), the only population with confirmed levamisole resistance in that study, whereas the variant was absent in all other populations analysed. The identification of Ser168Thr prompted us to look beyond *H. contortus*; a reanalysis of levamisole resistance in resequencing data from the closely related clade V parasitic nematode *Teladorsagia circumcincta* (*46*) revealed a homologous non-synonymous variant at high frequency in resistant parasites (Ser140Thr in Cont419:G75849C ; *freq*_Ser140Thr_ = 0.972), which was absent in the susceptible population to which it was compared. Although a serine to threonine substitution is a relatively conservative change, we found the serine residue to be highly conserved among clade V nematodes (**fig. S7**), particularly among the parasite species, whereas in the free-living *Caenorhabditis spp.*, threonine is encoded at this position.

In *C. elegans, acr-8* is genetically and functionally distinct relative to *acr-8* of parasitic nematodes and is not a component of the native levamisole receptor (*47*); the *C. elegans* functional homolog *lev-8*, which can be transgenically substituted by *H. contortus acr-8* to produce a functional receptor (*48*), does encode a serine at this homologous position. The *H. contortus* ACR-8 Ser168Thr variant lies immediately downstream of the cys-loop domain within the ligand-binding pocket and is immediately adjacent to a highly conserved tryptophan residue essential for ligand binding (*49, 50*) (**Fig. 3E**). Importantly, key residues downstream of the conserved tryptophan have previously been shown to influence levamisole sensitivity of closely related receptor subunits (*51*). Thus, we hypothesise that the Ser168Thr variant facilitates a change in the molecular interactions within the binding pocket of ACR-8, resulting in a decreased sensitivity to levamisole.

The identification of *lev-1* genes within the chromosome 4 QTL is compelling, with three intronic variants of *lev-1* (top variant position 14,995,062 in HCON_00107700; *P* = 1.7e-20; CMH test) among of the top ten most differentiated SNPs on this chromosome. However, it remains unclear what effect the overall observed variation in the *lev-1* genes has on levamisole resistance. Although multiple non-synonymous variants were also identified (seven and three variants for HCON_00107690 and HCON_00107700, respectively), none were predicted to cause high-effect changes in the protein sequence and exhibited only relatively minor shifts in allele frequency upon levamisole treatment. In *C. elegans*, several dominant resistant variants of *lev-1* have been described (not found in the data described here); however, *lev-1* can be lost without affecting the function of the receptor (*43*). Examination of variation in *lev-1* expression in addition to genetic variation may be required to elucidate the role of *H. contortus lev-1* subunits in levamisole resistance. Close to the *lev-1* genes and toward the centre of the QTL, four of the top ten variants in chromosome 4 were found in HCON_00107560 (top non-synonymous variant: Arg934His at position 14,781,344; *P* = 1.0e-21; CMH test), an ortholog of *C. elegans kdin-1*. Highly conserved with mammalian orthologs (*52*), *kdin-1* has been shown to co-localise with acetylcholine receptors at rat neuromuscular junctions during development (*53*) where, via a PDZ domain, it participates in the coordination of signalling components including ion channels and neurotransmitters. The precise role of HCON_00107560 or *kdin-1* in *H. contortus* or *C. elegans*, respectively, remains unknown; however, its putative association with levamisole response here warrants further investigation.

Signals of selection on two components of the pentameric acetylcholine receptor prompted us to look for selection on the remaining subunits. Although the expression of *unc-63* (HCON_00024380) and *unc-29.3* (HCON_00003520) mRNAs were significantly reduced in the larvae of resistant MHco18(UGA) strain (*54*), we found no evidence of selection on the region of the genome containing these genes.

### A resolved ivermectin QTL implicates cky-1 as a novel mediator of resistance

Ivermectin is a critically important broad-spectrum drug used to control several human- and veterinary-infective helminths worldwide and is also widely used as an acaricide targeting ticks and mites. We recently identified a ∼5 Mb QTL associated with ivermectin resistance from 37 to 42 Mb on chromosome 5 from the analysis of a backcross experiment (*34, 56*), and subsequently, we identified evidence of selection in the same chromosomal region in ivermectin-resistant field populations from Africa and Australia (*25*). Although the introgression region from the backcross was broad (*57*), the genetic architecture of the QTL was consistent with a single dominant variant driving resistance, and we were able to demonstrate that most candidate genes previously proposed to be associated with resistance were not under direct ivermectin selection. However, we were unable to confidently identify any novel candidate driving mutation among the ∼360 genes lying within the region (*34*).

Here, we confirm the QTL within the previously implicated chromosome 5 region at ∼37.5 Mb (*34, 56*) but with significantly increased resolution (**Fig. 4A**). We have narrowed the genetic association to approximately 300 kb wide (region: ∼37,250,000-37,550,000), based on the region of highest differentiation between independently replicated pre- and post-treatment X-QTL samples (**Fig. 4B**). This region was also highly differentiated between pre-treatment larvae and adult male worms that survived ivermectin treatment *in vivo* **(fig. S8 A,B),** and between larvae that survived treatment with an EC_75_ dose of ivermectin and those sensitive to an EC_50_ dose *in vitro* **(fig. S8 C,D)**. Together, these results confirm that this locus is under direct selection and mediates resistance in both the parasitic stages *in vivo* and in the free-living stages *in vitro* (see **Supplementary materials** for further discussion). Finally, this was the only region in the genome where increased levels of ivermectin resistance (i.e., EC_50_) was associated with a loss of genetic diversity in moderately or highly resistant field populations relative to susceptible populations (**Fig 4C**), consistent with a selective sweep in response to ivermectin-mediated selection.

**Fig. 4.**
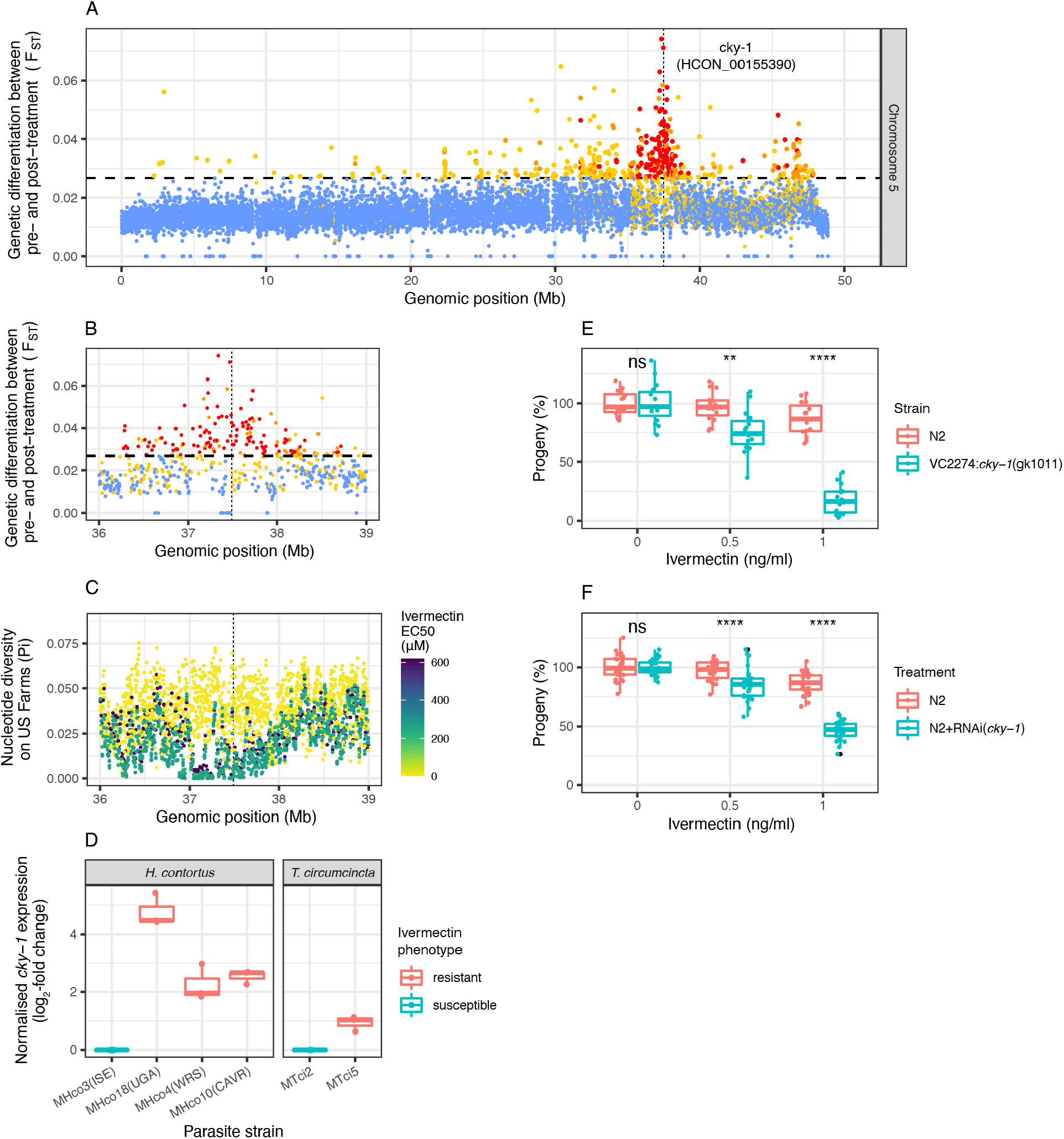
Characterisation of QTL associated with ivermectin resistance. (**A**) QTL between pre- and post-ivermectin treatment on chromosome 5. Each point represents the mean *F*ST in a five kb window; points are coloured based on the concordance of individual replicates indicated by none (blue), 1 of 3 (yellow), 2 of 3 (orange), or all 3 (red) above the genome-wide threshold (horizontal dashed line; mean + 3 SD of the chromosome-wide *F*ST). A magnified aspect of the main chromosome 5 QTL, highlighting (**B**) genetic differentiation (*F*ST) in the X-QTL cross, and (**C**) nucleotide diversity (Pi) on US farms, where each farm is coloured by the degree of ivermectin resistance (EC50) measured by larval development assays. In A, B and C, the position of *cky-1* is indicated by the vertical dashed line. (**D**) RT-qPCR analysis of *cky-1* expression in both *H. contortus* and *T. circumcincta* strains that differ in their ivermectin resistance phenotype. Data represents log2-transformed expression normalised to actin or GAPDH control genes for *H. contortus* and *T. circumcincta,* respectively. Downregulation of *cky-1* expression in *C. elegans* by either (**E**) a balanced deletion or (**F**) RNAi-knockdown increases ivermectin sensitivity relative to the control N2 strain, based on developmental assays measuring the percentage of progeny surviving to adulthood relative to DMSO controls. (In **E** and **F,** each point represents an independent treatment condition, which is normalised to a DMSO control without ivermectin. A Kruskal-Wallis test was used to determine whether treatment condition differed from untreated control, where ns = not significant, * *p* < 0.05, ** *p* < 0.01, and **** *p* < 0.0001.

The main chromosome 5 QTL contained 25 genes and included an expansion of protein kinases (8/21 genes present in the genome with the InterPro identifier IPR015897), some of which had the highest statistical association with resistance; for example, HCON_00155240 (intronic position 37,336,132, *P* = 3.3e-13; position 37,235,944, *P* = 1.2e-12) and HCON_00155270 (intronic position 37,343,439, *P* = 1.0e- 10). These protein kinases are, however, novel leads with no previous association to drug resistance, and a lack of functional orthologs and observed gene expansion made it difficult to further infer and test a role for these genes in ivermectin resistance.

Towards the middle of the QTL, we identified *cky-1* (HCON_00155390; positions 37,487,982 - 37,497,398) as a new mediator of resistance, based on several lines of evidence. In the X-QTL data, *cky-1* contained eight moderately to highly differentiated significant non-synonymous variants (top variant: position 37,497,061 [Ser583Pro], *P* = 9.6e-09; CMH test). In a complementary study, we showed *cky-1* was the only gene in the region significantly upregulated in both males and females of the resistant MHco18(UGA) isolate relative to MHco3(ISE) and was one of the most upregulated genes genome-wide (*58*). In this study, RT-qPCR of *cky-1* from the parental isolates of the cross and two unrelated ivermectin-resistant *H. contortus* strains revealed significant overexpression in ivermectin-resistant relative to sensitive strains (**Fig. 4E**), an observation that was replicated between sensitive and ivermectin-resistant strains of the related parasite, *T. circumcincta* (**Fig. 4E**). To explore *cky-1* further, we assayed *C. elegans* developmental and pumping behavioural phenotypes, both known to be perturbed by ivermectin exposure (*59*), to test the role of differential expression of *cky-1* on the resistant phenotype in the presence of ivermectin. While complete knockout of *cky-1* was non-viable, both a balanced deletion (VC2274) (**Fig. 4E**) and RNAi knockdown (**Fig. 4F**) of *cky-1* increased the sensitivity of *C. elegans* to ivermectin relative to the ivermectin-susceptible N2 strain. The level of *cky-1* expression is, therefore, associated with the ivermectin resistance phenotype in nematodes.

Two additional, less prominent QTLs on chromosome 5 at ∼46 Mb and on chromosome 2 at ∼3 Mb were also identified after ivermectin treatment (**Fig. 4A**; see **Supplementary materials** for a description of the two QTLs). The second chromosome 5 QTL was identified as a candidate region associated with resistance in the backcross (*34*); however, we did not have the statistical power to differentiate it from the main QTL in that experiment. Here, the QTL appeared to segregate independently of the prominent 37.5 Mb peak, providing more robust evidence of a second resistance-conferring variant on chromosome 5. Although the main chromosome 5 QTL at 37.5 Mb was present in all selection experiments with ivermectin, the secondary QTLs were variable between replicates and experiments. To further refine the association, we exposed the F5 generation of the cross to a half standard dose of ivermectin, followed by a double dose thereafter. The rationale was first to identify low-effect variants (responding to the half dose treatment), then select a subset of variants that conferred resistance at high doses (see **Supplementary materials** for additional background). In these experiments, we consistently detected the main chromosome 5 QTL but not the less prominent chromosome 2 QTL.

Additionally, we detected the presence of at least three new minor QTLs (**fig. S10**, supplementary text). Of practical significance, the identification of novel replicate- specific variants in addition to the main chromosome 5 QTL highlights the consequence of under-dosing in selecting novel variants, and emphasises the importance of correct dosing in the field.

## Discussion

Anthelmintics are currently the most important tool for controlling parasitic worm infections in humans and animals worldwide, and this is likely to remain true for the foreseeable future. However, this paradigm of control is threatened by the emergence and spread of anthelmintic-resistant parasites. Despite the large health and economic impacts resulting from increasing levels of anthelmintic resistance, multiple complicating factors have hindered the ability to determine the genetic loci responsible for resistance. Here we demonstrate an efficient approach to map multiple drug resistance-conferring loci for three of the most important anthelmintic drugs in the globally distributed and genetically tractable parasitic nematode, *H. contortus*. We have identified novel variants and loci likely involved in resistance to each of these drug classes; these include the β-tubulin isotype 2 Glu198Val variant correlated with benzimidazole resistance in field populations, the *acr-8* Ser168Thr variant associated with levamisole resistance in both the cross and field populations of *H. contortus*, and *cky-1* as a novel candidate gene that mediates ivermectin response. Our approach was validated by identifying QTLs and variants previously associated with drug resistance, for example, the β-tubulin isotype 1 Phe200Tyr variant associated with benzimidazole resistance and the *acr-8* indel variant associated with levamisole resistance. However, for the latter, we provide evidence against the indel being a reliable marker of resistance. Finally, we note an absence of many previously proposed ivermectin- associated candidate genes in the QTL described, highlighting both the limitation of candidate gene approaches and the power of genome-wide forward-genetic strategies to robustly identify regions of the genome containing known and novel mediators of resistance (*9*).

We have refined a previously identified QTL for ivermectin resistance on chromosome 5 (*34*) to ∼300 kb, and together with functional genetic evidence from expression and knockout experiments, we have explicitly tested the role of our proposed candidate in the main ivermectin QTL on chromosome 5, the NPAS4 ortholog *cky-1*. This gene encodes an activity-dependent basic Helix-Loop-Helix (bHLH)-PAS family transcription factor shown in mammals to regulate the excitation/inhibition balance upon neuronal activation to limit excitotoxicity (*60*) and during the development of inhibitory synapses to control the expression of activity-dependent genes (*61*). It is yet to be determined if this is a conserved molecular function in nematodes; however, it is tempting to speculate that the hyperexcitability as a result of induced activation of ion channels by ivermectin at the neuromuscular junction is, at least in part, controlled by a “retuning” of the excitation/inhibition balance to limit toxicity. The role of *cky-1* in ivermectin resistance is supported by: (i) genetic differentiation between susceptible and resistant strains around this locus relative to genome-wide variation that is replicated in geographically and genetically diverse strains here and elsewhere (*25, 34, 62*), (ii) the presence of non-synonymous variants that are highly differentiated before and after treatment, (iii) increased gene expression of *cky-1* in resistant strains relative to susceptible strains (supported by genome-wide RNA-seq (*58*)) and (iv) knockdown of the *C. elegans* ortholog leading to hypersensitivity to ivermectin. We acknowledge that overexpression of *cky-1* in *C. elegans* does not recapitulate the high levels of ivermectin resistance seen in *H. contortus* or, for example, by concurrent mutation of glutamate-gated chloride channels in *C. elegans* (*21*); while this may argue against *cky-1* as a universal mediator of resistance, it likely reflects the challenge of using a heterologous expression system in which there is an assumption that the biology (and, therefore, response to treatment) is concordant between the free-living and parasitic species, and/or may reflect the multigenic nature of ivermectin resistance in different species (*63–65*). Given the lack of an obvious causal non-synonymous variant, we hypothesise that a non-coding variant that influences the expression of *cky-1* is under selection in resistant strains of *H. contortus*; however, such variants are difficult to validate without genotype and transcriptional phenotype data from a large number of individual worms.

It is broadly accepted that the mode of action of ivermectin is on ligand-gated ion channels, and ivermectin resistance has been associated with variants in glutamate-gated channels (*66*). Concurrent mutation of a number of these channels (*glc-1, avr-14* and *avr-15*) confers high-level resistance in *C. elegans* (*21*) and selection on at least one of these channels (*glc-1*) in wild strains (*67*) has been demonstrated. We find no evidence to suggest that genetic variation in these channels confers ivermectin resistance in *H. contortus*. Transcriptional changes in these channels in resistant, relative to drug-susceptible, parasite strains have been demonstrated previously; for example, the glutamate-gated chloride channel subunits (*glc-3, glc-5*), as well as p-glycoprotein ABC transporters (*pgp-1, pgp-2, pgp-9*) (*54*) in the MHco18(UGA) strain. Similarly, a *pgp-9* copy number variant was associated with ivermectin resistance in a genetic cross and bulk segregant experiment in the related nematode *T. circumcincta* (*46*), while transgenic overexpression of the equine parasitic nematode *Parascaris univalens pgp-9* modulated ivermectin sensitivity in *C. elegans* (*68*). However, none of these genes were identified in regions of differentiation after treatment in this study, suggesting these genes are not the direct target of selection. However, we cannot exclude that variation in expression of these genes may be a downstream response to selection on a transcriptional regulator such as *cky-1*.

The use of genetic crosses, in which the genetics of the parasites can be controlled, is the ideal way to generate populations of individuals in which the relationship between genotype and phenotype can be assayed. Our approach here relies on selecting populations of parasites using drug treatment, however, advances are still required to improve phenotyping of resistance in individual parasites. The ability to do so would improve our understanding of the molecular basis of drug resistance phenotypes and enable more sophisticated genetic approaches to unravel the role of the minor signatures of selection we observe in this experiment. Recent advances in single larvae whole-genome sequencing (*69*) and low-input RNA sequencing (*70*), even at single-cell resolution (*71*), now provide the tools to allow a more precise understanding of molecular and cellular phenotypes for drug response and may help to fully understand the role of *cky-1*. The identification of *cky-1* as a putative candidate offers new plausible hypotheses relevant to a resistant phenotype, whereby *cky-1* may act: (i) during development to establish a neuronal architecture that is more tolerant to hyperexcitability such as that caused by ivermectin, and/or (ii) in response to ivermectin exposure by initiating transcription of downstream genes to modulate the excessive excitation/inhibition imbalance, thereby mitigating the lethal effect. These hypotheses will require further validation, aided in the first instance by identifying the downstream targets of *cky-1*. However, it is clear that the molecular mechanisms by which parasites develop ivermectin resistance are more complex than previously appreciated. Broader, systems biology approaches are likely needed to understand the relationship between direct evidence of selection in the genome and the downstream transcriptional responses that enable parasite survival when challenged with ivermectin. By defining the genomic landscape of anthelmintic resistance even in a single resistant strain, our results refocus effort away from candidate genes with limited support and redefine our understanding of the evolution of anthelmintic resistance in helminths of veterinary and medical importance.

## Acknowledgements

This work was supported by the Biotechnology and Biological Sciences Research Council (BBSRC) [BB/M003949]; the Scottish Government’s Rural and Environment Science and Analytical Services (RESAS) division; a University of Glasgow James Herriot Scholarship; the Wellcome Trust [206194 and 216614/Z/19/Z]; and UKRI [MR/T020733/1]. For the purpose of Open Access, the authors have applied a CC BY public copyright licence to any Author Accepted Manuscript version arising from this submission. We would like to acknowledge members of the BUG Consortium and Parasite Genomics group at the Wellcome Sanger Institute for insightful discussions throughout this project. We also thank Pathogen Informatics and DNA Pipelines (WSI) for their support and expertise and the Biosciences Division at the Moredun Research Institute for expert care and animal assistance.

## Author contributions

Conceptualisation: RL, AT, ED, JAC. Investigation: SRD, RL, DB, AM, KM, AA, CB, UC, IF, JM. Formal analysis: SRD, RL. Software: SRD. Resources: DB, AM, RK, NS. Supervision: CB, JSG, NS, MB, ED, JAC. Project administration: NH, ED. Writing - Original Draft: SRD, JAC, RL, ED. Writing - Review & Editing: All authors. Funding acquisition: RL, AT, MB, ED, JAC, NS, BM, DB, CB, SRD.

## Data and materials availability

Raw sequencing data for this study are outlined in **table S1** and are archived under the ENA study accession PRJEB4207. The *H. contortus* genome assembly and manually curated annotation resources are publicly available at https://parasite.wormbase.org/Haemonchus_contortus_prjeb506/Info/Index/. The code used to generate and analyse data and to plot figures can be found at https://github.com/stephenrdoyle/hcontortus_X-QTL.

## Supplementary Materials

Materials and Methods

Supplementary Text

Figs. S1 to S10

Tables S1 to S2

References (72–97)

