## Supplementary Materials for "Genomic landscape of drug response reveals novel mediators of anthelmintic resistance"

**This PDF file includes:**

Materials and Methods
Supplementary Text
Figs. S1 to S10

**Other Supplementary Materials for this manuscript include the following:**

Tables S1 to S2

### Materials and Methods

Animal handling and ethics statement

All experimental procedures were examined and approved by the Moredun Research Institute Experiments and Ethics Committee and were conducted under approved UK Home Office licenses following the Animals (Scientific Procedures) Act of 1986. The Home Office licence number is PPL 60/03899, and the experimental code identifier is E46/11.

Establishment of the genetic cross between a susceptible and multi-drug resistant strain of *H. contortus*

This study extends the analysis of a genetic cross initially described by Doyle and colleagues (*57*) between the anthelmintic susceptible strain MHco3(ISE) (*72*) and MHco18(UGA), a field derived strain of *H. contortus* that is insensitive to standard treatment doses of benzimidazoles, levamisole, and ivermectin (*54*). Briefly, two sheep were orally infected with *H. contortus* third-stage infective larvae ($L_3$); one with 10,000 $L_3$ of the MHco3(ISE) isolate and one with 10,000 $L_3$ of the MHco18(UGA) isolate. Immature adult worms were recovered from both sheep at necropsy on day 14 post-infection. After being rinsed in physiological saline at 37°C and sexed based on morphology, 100 female MHco3(ISE) and 100 male MHco18(UGA) worms were surgically transferred into the abomasum of a third sheep. Faeces were collected from day three post-transfer, F1 eggs were coprocultured for 14 days to develop, after which the $L_3$ larvae were retrieved by the Baermann technique. A fourth sheep was orally infected with ~5,000 F1 $L_3$, and from day 21 post-infection, we collected F2 eggs and cultured them to $L_3$ as above. For all generations of the genetic cross, the $L_3$ were either maintained in tap water at 8°C or exsheathed and snap-frozen in liquid nitrogen and stored for future infections. Adult worms were recovered at necropsy, rinsed, sexed and snap-frozen in batches of 20 males or females before archiving at -180 °C.

Anthelmintic selection on the F2 generation with three anthelmintics (X-QTL)

The core aim of the work described was to perform a drug selection experiment on adults from the F2 generation of the genetic cross, followed by X-QTL genetic mapping of drug-specific genetic loci on F3 progeny by whole genome sequencing and analysis. The experiments were performed in triplicate. Briefly, F2 larvae were used for oral infection of 12 donors, and eggs were collected from day 21 and cultured to $L_3$. On day 35, donor sheep were treated with either (i) 0.2 mg/kg ivermectin (Oramec Drench, Boehringer Ingelheim Animal Health, UK), (ii) 7.5 mg/kg fenbendazole (Panacur 10% Oral Suspension, MSD Animal Health), (iii) 7.5 mg/kg levamisole hydrochloride (Levacide Low Volume, Norbrook), or (iv) left untreated as a control. Eggs were collected from all donors for 21 days post-treatment and cultured to $L_3$ as described above. Larvae collected pre- and post-treatment and from time-matched controls were snap-frozen in batches of 200 $L_3$ for DNA extraction. A follow-up experiment to improve the resolution of QTLs by increasing the recombinants per pool was performed; in this case, ~5000 $L_3$ were pooled for DNA extraction. On day 56 post-infection, all donors were euthanised, and adult worms were harvested and stored as described above.

F3 selection with ivermectin (Advanced Intercross)

We performed subsequent selection experiments to refine the QTL from the primary X-QTL analyses. The advanced intercross experiment, which involved drug treatment at half standard-dose followed by a subsequent double standard-dose treatment of ivermectin was performed on the F3 generation adults, after which the F4 progeny were collected for whole genome sequencing and analysis. The experiment was performed in triplicate. Briefly, pre-treatment F3 generation $L_3$ from three control donors in the X-QTL selection experiment were pooled, and aliquots of ~5,000 $L_3$ were used to infect seven donors (four for IVM treatment, including one ‘test’ donor to ensure adult parasites survived the treatment regime, and three untreated controls). Eggs were collected from day 21. On day 28, four donors were treated with 0.1 mg/kg (half standard dose) ivermectin (Oramec Drench, Boehringer Ingelheim Animal Health, UK), and eggs were collected from all donors for the next seven days. On day 35, the test donor was treated with 0.4 mg/kg (double standard dose) ivermectin and continued to produce eggs over the following seven days. The remaining three donors on the drug treatment regime were given 0.4 mg/kg ivermectin on day 42. Eggs were collected for 14 days post-treatment from all donors. Eggs produced pre- and post-ivermectin treatment and from time-matched untreated controls were cultured to $L_3$ and snap-frozen in batches of 200 larvae. On day 56, all donors were euthanised, and adult worms were harvested and stored as described above.

*In vitro* larval development and dose-response assays

We used the commercial DrenchRite larval development assay to quantify the effective concentration of anthelmintic that resulted in various proportions of the maximum effect of the drug (i.e. $EC_{50}$ is the concentration to produce 50% of the maximum). The basis for the assay is the observation that eggs from drug-resistant populations hatch in the presence of ivermectin, levamisole or thiabendazole and develop to $L_3$ over 6-7 days. In contrast, eggs from drug-sensitive populations fail to hatch (levamisole and thiabendazole) or hatch then arrest at $L_1$ (ivermectin). The proportion of eggs that develop to $L_3$ across a range of increasing drug concentrations is used to generate a dose-response curve, from which the $EC_{50}$ is used as a measure of the resistance status of the adult population (*73*).

We used this approach to:

1) determine the $EC_{50}$ for the parental isolates and F3 generation of the genetic cross, in order to quantify the resistance status of the parental and admixed parasites; and

2) determine the $EC_{25}$, $EC_{50}$ and $EC_{75}$ for the F5 generation of the genetic cross in order to phenotypically define drug-susceptible and highly-resistant larvae for QTL mapping.

The standard DrenchRite assay was performed as follows. Briefly, eggs were isolated from fresh faeces using standard procedures and diluted to 10,000 eggs in 2 ml tap water with 22.5 µg/ml amphotericin B (Thermo Fisher, 15290018). The egg suspension was vortexed thoroughly, and 20 µl (~100 eggs) was added to each well of a 96-well DrenchRite plate (*73*). Plates were sealed with parafilm and incubated at 26°C. After 24 h, when most eggs had hatched, 20 µl nutritive media (*73*) was added per well. On day 6, the assay was terminated with Lugol's iodine stain (Fisher Scientific, 12996307), and the contents of each well were transferred to flat-bottomed 96-well plates. The numbers of eggs, $L_1$, $L_2$ and $L_3$ were counted per well using an inverted light microscope at 100-200× magnification. A Probit model was fitted to identify the $EC_{25}$, $EC_{50}$ and $EC_{75}$ for ivermectin.

Next, to isolate large numbers of sensitive $L_1$ at a low concentration of ivermectin and resistant $L_3$ developing normally in the high concentration ivermectin conditions for QTL mapping, we developed an in-house "scaled-up" larval development assay for the F5 generation. To perform the scaled-up larval development assay, ivermectin aglycone (Santa Cruz Biotechnology, CAS 123997-59-1) was diluted in DMSO and added to 2% Nematode Growth Medium (NGM) agar without cholesterol to give final ivermectin concentrations that corresponded to those used in the DrenchRite assay: 3.9 nM ($EC_{25}$), 15.6 nM ($EC_{50}$) and 62.5 nM ($EC_{75}$). The assay was performed in 12-well plates and triplicate wells were prepared for each ivermectin concentration and DMSO controls. The experimental conditions of the scaled-up assay were as described above, except that 1000 eggs were added per well in 200 µl of tap water with amphotericin B, 200 µl nutritive media was added after 24 h, and the assay was stopped on day 5 (without iodine fixation) to retrieve sensitive $L_1$ before degradation.

In our initial DrenchRite assays, the DMSO control wells typically showed between 75 and 95% of the population developing to $L_3$, with a small proportion (5-25%) that would hatch, but not develop normally. We reasoned that the $L_1$ in the $EC_{25}$ wells would also likely include a proportion of these hatched but non-viable larvae; given we were trying to isolate truly susceptible parasites, it was important to distinguish these non-viable larvae from drug-susceptible larvae. However, because larvae could not be fixed before sequencing and high power microscopy was impractical at this scale, we could not undertake full counts to measure development accurately. For this reason, sensitive $L_1$ were isolated from the $EC_{50}$ wells (defined as arrested $L_1$ stages), and resistant $L_3$ were isolated from the $EC_{75}$ wells to separate the population for sequencing.

Sampling and dose-response of parasites from US farms

To complement the selection experiments derived from the genetic cross and X-QTL analyses, we sampled pools of $L_3$ from nine farm populations in the US. These samples were collected as part of routine anthelmintic resistance screening and phenotyped for ivermectin, levamisole and benzimidazole resistance using the DrenchRite assay as described above (*74*). See **table S2** for $EC_{50}$ data for the three drug classes. These farms have applied different management strategies and drug exposure histories and thus the worms will have been exposed to different drug selection pressure(s). While we do not have complete detail of the management history of these populations, we selected populations for comparative analysis from a larger collection of farms based on these DrenchRite $EC_{50}$ data, from which an ivermectin susceptible, three moderately resistant and five highly resistant farm populations to ivermectin were chosen (see **fig. S2** and **S3**). Due to the limited number of fully drug-sensitive populations from commercial farms, one additional drug-sensitive isolate from the University of Georgia was included (UGA-SUSC (*75*); Farm 1). The populations selected also have variable levels of both benzimidazole and levamisole resistance; therefore, we used these populations to validate the candidate regions associated with all three drug classes that are likely to be under drug selection in the field.

Larvae were archived at -80°C in water or 70% ethanol, then thawed, rinsed in PBS and snap-frozen in batches of 200 $L_3$ prior to DNA extraction.

Sample preparation and whole-genome sequencing

Genomic DNA was isolated from pools of 200 $L_3$ or individual adult males as follows: 20 µl of 20 mg/µl proteinase K (ThermoFisher Scientific, 25530031) together with 300 µl lysis buffer (200 mM NaCl, 100 mM Tris-HCl, 30 mM EDTA pH 8, 0.5% SDS) was added to the frozen pellets of larvae or adult worms before incubation at 55°C for 2 h. Next, 10 µl of 10 mg/ml RNAse A (ThermoFisher Scientific, EN0531) was added before incubation at 37°C for 10 min. 550 µl phenol/chloroform/isoamyl alcohol (25:24:1) (ThermoFisher Scientific, 15593031) was added to the lysate, shaken vigorously for 15 s, incubated at room temperature for 5 min, then centrifuged at 14,000 g for 15 min at room temperature. The top layer was carefully removed to a fresh tube, and 0.1× volume sodium acetate pH 5.5 was added, followed by 3× volume 100% EtOH at room temperature, then 2 µl glycogen, before overnight incubation at -80oC. After 5 min of centrifugation at 14,000 g at 4oC, the supernatant was carefully aspirated, and 500 µl 70% EtOH was added to the pellet before centrifugation for another 5 min at 14,000 g at 4oC. The supernatant was carefully removed over a lightbox to visualise the pellet before a brief spin to facilitate aspiration of any remaining EtOH. The pellet was airdried until all traces of EtOH were removed before re-suspending in ≥10 µl EB buffer (Qiagen, 19086). Genomic DNA was stored at 4oC.

Individual sequencing libraries (mean length of ~400 bp) were prepared using a PCR-free protocol as previously described (*76*) and sequenced on Illumina HiSeq2500 (X-QTL & advanced intercross) and HiSeq4000 (US farm) platforms using 125 bp and 150 bp PE chemistry, respectively. We aimed to perform sufficient sequencing to achieve 100-fold coverage of the ~283 Mb genome for each sample sequenced.

Mapping and variant calling

Raw sequence data were first assessed for quality using Fastqc (http://www.bioinformatics.babraham.ac.uk/projects/fastqc/) and visualised using MultiQC (*77*). Reads were first trimmed using trimmomatic v0.32 (*78*) before they were mapped to the *H. contortus* V4 reference genome (*33*) (available here: https://parasite.wormbase.org/Haemonchus_contortus_prjeb506/Info/Index/) using BWA-MEM (bwa: v0.7.12-r1039; (*79*))(parameters: -Y -C -M). Duplicate reads were marked using Picard (v2.5.0; https://github.com/broadinstitute/picard), after which perfectly mapped paired reads were extracted using Samtools-1.3 view (-f 14). Finally, indel realignment was performed using GATK v3.7.0 IndelRealigner (*80*). Genetic variation was determined using Samtools-1.3 mpileup (parameters: -F 0.25 -d 500 -b bamlist), which was performed for each experimental group of samples separately (i.e. each drug treatment group including all replicates). To determine the predicted functional consequence of individual variants, mpileup files were processed using bcftools v1.9 (parameters: call -vm -Oz), after which annotation of the variant call format (VCF) file was performed using SNPeff (*81*) (parameters: -no-intergenic -no-downstream -no-upstream). The current annotation GFF used to build the annotation database is available at WormBase ParaSite (release 15; https://parasite.wormbase.org/index.html).

Genome-wide analyses of genetic variation

To determine within sample measures of nucleotide diversity, multi-sample mpileup files were first processed to generate a single pileup file per sample per chromosome, which were subsequently used as input to npstat (parameters: -n 400 -l 5000 -mincov 20 -maxcov 200 -minqual 20 -nolowfreq 2)(*82*).

Between-sample measures of genetic diversity were determined using Popoolation2 (*83*); briefly, a synchronised file was first generated from each mpileup (popoolation2 mpileup2sync.jar --min-qual 20). Pairwise estimates of genetic differentiation were first determined by $F_{ST}$ calculated in 5 kbp windows throughout the genome (popoolation2 fst-sliding.pl --pool-size 200 --window-size 1000 --step-size 500 --min-count 4 --min-coverage 50 --max-coverage 2%), and secondly using a Fisher's exact test on individual variants (popoolation2 fst-sliding.pl --window-size 5000 --step-size 5000 --min-count 4 --min-coverage 30 --max-coverage 2%). Finally, a Cochran-Mantel-Haenszel (CMH) test of independence was undertaken to determine concordance of replicate samples per variable site (popoolation2 cmh-test.pl --min-count 2 --min-coverage 30 --max-coverage 2% --population).

Analyses of specific variants identified in the genome-wide analyses, including calculation of allele frequencies and extraction of annotated SNPs based on functional effect classes, were performed using vcftools v0.1.16 (*84*).

Analyses of *acr-8* variation

We explored variation in *acr-8*, a gene previously implicated in levamisole resistance and a candidate that we identified in our X-QTL analyses of levamisole selection, in a number of different ways. First, we calculated the frequency of the *acr-8* indel, a marker previously associated with resistance, by counting sequencing reads containing the deletion pre- to post-treatment based on sequencing read coverage within and outside the indel for each replicate condition. Second, upon the identification of the Ser168Thr variant as being highly associated with resistance, we compared the protein sequence conservation of ACR-8 among clade V nematodes by aligning protein sequences obtained from WormBase Parasite using mafft v7.407 (*85*), which were visualised using the ggmsa (https://cran.r-project.org/web/packages/ggmsa/vignettes/ggmsa.html) R package. Finally, we explored the putative protein structure of ACR-8, again with particular reference to the position of the candidate levamisole-resistance associated variant Ser168Thr, using the I-TASSER (Iterative Threading ASSEmbly Refinement) webserver (https://zhanglab.dcmb.med.umich.edu/I-TASSER/) (*86*) and compared predictions against known molecular structures at the Protein Data Bank.

Real-time quantitative PCR (RT-qPCR) of *cky-1*

In order to more explicitly investigate functional changes at the *cky-1* locus, its expression was investigated in ivermectin susceptible and resistant *H. contortus* isolates. Total RNA was extracted from 20 male worms from each of three different donor sheep per isolate (MHco3[ISE]), MHco18[UGA], MHco4[WRS] and MHco10[CAVR]). 1 µg RNA was used for oligo(dT) primed cDNA synthesis (SuperScript® III First-Strand Synthesis System, ThermoFisher, 18080051), with no-reverse transcriptase controls included for each sample. cDNA was diluted 1:10 for RT-qPCR and 1 µl added to each reaction. RT-qPCR was undertaken on triplicate samples following the Brilliant III Ultra Fast SYBR QPCR Master Mix protocol (Agilent Technologies, 600882) using the Mx30005P system (Agilent) and results were analysed with MxPro v4.10. Gene expression was normalised to β-actin (HCON_00135080). All primer sequences are listed in Table S1.

In order to test the ubiquity of the association with resistance, expression of *cky-1* was also assessed in *T. circumcincta*; the same methods were used, except total RNA was extracted from 20 male worms from the susceptible MTci2 and ivermectin resistant MTci5 *T. circumcincta* isolates, a 1:5 cDNA dilution was used, and gene expression was normalised to *gapdh-1* (*87*).

**Table S1.** Primer sequences for RT-qPCRs

| Primer name | Sequence |
|---|---|
| HCON_00135080_F1 | CCAGTTGGTGACGATTCC |
| HCON_00135080_R1 | GGGTTTGCTGGAGATGACG |
| HCON_00155390_F1 | CCGAGACCAGATCAATGTCG |
| HCON_00155390_R1 | CACACTGTCTTCGGCTATCG |
| TCIRC_GAPDH1_F | TTGAGAAACCAGCTAGCATGGA |
| TCIRC_GAPDH1_R | CGCACCCTCCGAAGCA |
| TCIRC_CKY1_F | TCAGCGATGGCAATGGAAG |
| TCIRC_CKY1_R | GAAGGACCGCCAAAAAGAG |

RNA interference (RNAi) of *cky-1* in *C. elegans*

To explore the effect of *cky-1* expression on ivermectin response, we hypothesised that RNAi knockdown of *cky-1* expression would increase the sensitivity of *C. elegans* to ivermectin. A 299 bp fragment of *Cel-cky-1* was amplified by PCR (Pfu Ultra II Phusion polymerase; see Table S2 for primers) from adult *C. elegans* cDNA and purified using a PCR clean up kit (Qiagen, 28104). This product was cloned into TOPO-TA 2.1 and Sanger sequenced (Eurofins) using T7 primers to confirm its identity and that the full-length sequence was present. It was then sub-cloned by SacI/XbaI restriction digest followed by ligation into the L4440 double T7 RNAi expression vector (L4440 was a gift from Andrew Fire; Addgene plasmid #1654; http://n2t.net/addgene:1654 ; RRID:Addgene_1654) and sequenced again to confirm the construct.

**Table S2.** Primer sequences to amplify and clone a *cky-1* fragment for RNAi

| Primer name | Sequence |
|---|---|
| Ce_CKY1_299bp_RNAi_F_Sac1 | CGGAGCTCTGCAGATGTCATCTCCAAT |
| Ce_CKY1_299bp_RNAi_R_Xba1 | GCTCTAGACTGTCGTTTTGCAGTTCTTGC |

Double-stranded RNA (dsRNA) was prepared from an L4440:Ce299bpCKY1 clone using the Ambion Megascript kit (ThermoFisher Megascript T7 transcription kit, AM 1333). The following mixes were prepared in 200 µl PCR thin-walled RNase- and DNase-free tubes: three tubes containing 2 µl of 5X soaking buffer (1.25X M9 $Mg^{2+}$ free, 15 mM spermidine (Sigma S2626), 0.25% gelatin) and 8 µl of dsRNA CeCKY1 299 bp fragment; three tubes containing 2 µl 5X soaking buffer and 8 µl of elution buffer from the dsRNAi Megaclear kit (Ambion AM1908).

*C. elegans* N2 (Bristol "wild-type" laboratory strain) worms were grown to early $L_4$ stage on standard Nematode Growth Medium (NGM) plates containing *E. coli* OP50. Prior to RNAi-treatment, individual worms were picked onto empty NGM plates (without added OP50) and allowed to crawl for a few minutes to remove any residual OP50. 30 $L_4$ were transferred into the prepared RNAi or control tubes and incubated at 20°C for 24 h. Drug plates were prepared with DMSO or ivermectin to a final concentration of 0.5, 1.0 and 2.0 ng/ml ivermectin; a single treated worm from either RNAi or buffer-only tubes was transferred to each of 10 plates for all drug concentrations. The plates containing worms were then placed at 20°C for four days and assessed for development. All worms that reached the $L_4$ stage and above were counted and recorded.

We also measured the effect of the reduced *cky-1* expression on ivermectin sensitivity using a balanced deletion line of *C. elegans*, VC2274 [cky-1(gk1011) V/nT1 [qIs51] (IV;V)]. Five $L_4$ worms were transferred onto each of five plates and left at 20°C for five days. Additional worms and a longer assay duration were used due to the lower fecundity and slower development rate to adulthood, respectively, of the VC2274 line relative to N2. The numbers of worms reaching $L_4$ stage and above were counted. Data presented are from three separate experiments.

### Data availability

Raw sequencing data for all experiments are described in Table S1 and are available under the European Nucleotide Archive (ENA) study accession PRJEB4207. The *H. contortus* genome assembly and manually curated annotation resources are publicly available at https://parasite.wormbase.org/Haemonchus_contortus_prjeb506/Info/Index/.

### Code availability

Custom code that was used to analyse data and produce the figures presented is available at https://github.com/stephenrdoyle/hcontortus_xqtl and Zenodo.

### Supplementary Text

<u>Establishment of a stable triple-resistant genetic cross</u>

We crossed females of the fully drug-susceptible MHco3(ISE) isolate of *H. contortus* with males of the triple-resistant MHco18(UGA) isolate *in vivo*. The success of the cross was confirmed by the production of eggs by MHco3(ISE) females representing the F1 generation, their development to viable infective ($L_3$) larvae (validated by whole genome sequencing; (*57*)), and the establishment of fertile adult F1 in a donor sheep. Larvae were passaged without drug selection to generate the F2 and all subsequent generations of the genetic cross.

To assess drug efficacy in the F2 adults used for X-QTL analysis, faecal egg counts were undertaken throughout the course of the infection. Although there were insufficient sheep to undertake a full faecal egg count reduction test or controlled efficacy test, faecal egg counts for the three sheep in each group were compared pre-treatment (on day 35) and 7 or 14 days post-treatment for benzimidazole/levamisole and ivermectin, respectively. Based on the reduction in egg output, the F2 generation contained adult parasites that were resistant to standard doses of all three drugs, confirming the establishment of a stable triple-resistant genetic cross. However, the resistance level differed greatly; benzimidazole and levamisole showed 73% and 94% efficacy, respectively, while ivermectin showed 10% efficacy. These findings are broadly consistent with the predicted mode of inheritance of resistance to each drug (partially recessive for benzimidazole and levamisole (*92*), dominant or partially dominant for ivermectin (*34*, *93*).

<u>Genetic structure of the benzimidazole QTL</u>

After selection with benzimidazole, a single broad QTL was found on chromosome 1 which contains the β-tubulin isotype 1 gene, a locus with previously described variants that confer benzimidazole resistance. However, we note that Phe200Tyr was not the most significantly differentiated variant between pre- and post-treatment in the region; 817 polymorphic sites including eight non-synonymous mutations were identified on chromosome 1 with a *P*-value equal to or lower than Phe200Tyr. This highlights one of the challenges of identifying causal mutations in a highly genetically diverse species even under the controlled conditions of the genetic cross.

The β-tubulin isotype 1 gene was not in the middle of the QTL as expected, but was found on the right edge of an approximate 3 Mb region of differentiation extending downstream. The relevance of this extended genetic structure within the QTL is unclear. Although a QTL surrounding the β-tubulin isotype 1 locus was present in the US farms that are phenotypically resistant to benzimidazoles (**fig. S3**), the additional downstream region of differentiation was absent. The broader QTL from the genetic cross may reflect an artefact of the cross due to linkage disequilibrium with the Phe200Tyr mutation; however, the high recombination rate in this region in this cross (*57*) suggests that this is unlikely. Alternatively, this broader region may represent selection on additional, novel loci that require further investigation. For example, it is notable that two of the eight non-synonymous variants were found in cathepsin B-like cysteine proteinases (HCON_00005790 & HCON_00005960), which have previously been implicated in benzimidazole-mediated damage of intestinal cells of *H. contortus* (*94*, *95*).

Ivermectin resistance mediated by the chromosome 5 QTL is consistent between indirect and direct measures of selection

The X-QTL analyses presented were performed on the progeny (environmentally accessible eggs developed to $L_3$ in faecal cultures) of adult parasites directly exposed to anthelmintics *in vivo*. Therefore, measures of selection on genetic diversity were indirect; as the progeny sampled are not exposed to drug treatment, this approach assumes that the genetic variation sampled in the progeny accurately reflects the genetic variation in the adults that are directly exposed to drug treatment *in vivo*. Since the mode of inheritance of ivermectin resistance is likely dominant or semi-dominant (based on previous work in different *H. contortus* strains that share the same chromosome 5 QTL where F1 individuals are ivermectin resistant (*34*, *93*)), mating between phenotypically-resistant heterozygous adults will produce ~25% phenotypically susceptible, homozygous progeny. The consequence of this is a reduction in both: (i) the overall frequency of the resistant allele in the post-treatment progeny populations; and (ii) the genetic differentiation between pre- and post-treatment populations (i.e. the causative resistant allele will not go to fixation in the surviving, post-treatment progeny population due to the presence of phenotypically susceptible individuals). To address this, we performed two additional complementary experiments to test the effect of direct selection. First, we sequenced pools of F3 adult male worms that survived treatment and compared them against pooled control $L_3$ from the same generation. The genome-wide genetic differentiation between the two groups was generally more variable (**fig. S8 A**), perhaps due to the lower number of individuals sampled (n = 40 adult males), the comparison between single-sex adults and mixed-sex $L_3$, and/or comparisons between populations sampled from different hosts (as opposed to the X-QTL experiments in which the pre- and post-treatment were sampled from the same host). However, the chromosome 5 QTL was still discernible with the peak of differentiation between 37.3 and 37.5 Mbp (**fig. S8 B**). Second, we performed *in vitro* larval development assays on the F5 generation of the cross, where eggs were cultured to $L_1$/$L_2$ or $L_3$ in a concentration gradient of ivermectin. We first performed a dose-response experiment to determine the $EC_{25}$, $EC_{50}$ and $EC_{75}$ of the population with the DrenchRite assay (**fig. S8 C**). This experiment was followed by a larger-scale development assay at relevant concentrations to isolate large numbers of larvae for whole-genome sequencing. Two groups of larvae were obtained and analysed: (i) those that were susceptible to low concentrations of ivermectin (15.6 nM and below) and arrested at the $L_1$ stage, and (ii) those that were resistant to ivermectin and proceeded to develop at high concentrations (62.5 nM and above). Again, a clear QTL at the chromosome 5 locus with comparatively little background variation was observed between the two groups (**fig. S8 D**). These additional experiments both provide evidence to support the importance of the major chromosome 5 QTL in ivermectin resistance and that this signal of genetic differentiation is consistent in assays where selection is applied either directly or indirectly following drug exposure.

### Recombination and its impact on the resolution of Pool-seq QTL

The resolution to map causal loci using a genetic cross and X-QTL approach is somewhat dependent on the amount of recombination that has taken place and thus is a function of the number of recombinant individuals present. We assayed variation from 200 progeny per pool, a number somewhat arbitrarily chosen but one that we thought would allow sufficient recombinants and reproducibility between replicate experiments; as a consequence, this in part limited our ability to refine the QTL regions to relatively broad regions of genetic differentiation. Our ability to further refine these QTL to identify causal variants was further confounded by high levels of background (non-causal) genetic variation within the QTL that is linked to the causal variant and co-inherited from the founding resistant strain. Increased resolution could be achieved by increasing the sample size per pool (see **fig. S8** for preliminary data using ~5000 $L_3$ progeny per pool), as has been recently demonstrated using *C. elegans* in which millions of individuals were used per pool (17). Further resolution towards defining causal loci could be achieved by reducing the genetic variation in the founding cross, either by crossing phenotypically distinct parental strains that are genetically more closely related, or by experimental evolution whereby a resistance phenotype is gradually selected for from a susceptible population using increasing drug exposure over time.

### Characterisation of the minor loci associated with ivermectin resistance on chromosomes 2 and 5

The minor QTL on chromosome 5 (range: 45,000,000-47,500,000) was broad and contained 221 genes, including two genes *avr-15* (HCON_00161180; 45,134,921-45,149,462) and *pgp-11* (HCON_00162780; 47,243,601-47,259,265) that have been associated with ivermectin resistance previously (*59*, *96*). Here, both genes were found on the boundaries of the QTL, and while they do contain statistically significant non-synonymous variants (three and five for *avr-15* and *pgp-11*, respectively), the differences in variant frequency between pre- and post-treatment are relatively minor compared with other variants within the region leading us to conclude that they are unlikely to be the direct target of selection here. Finally, the minor QTL towards the start of chromosome 2 (region: 2,992,500-3,267,500) contained two previously described candidate genes - *osm-1* (HCON_00035760) and *che-11* (HCON_00035880) - that have been shown to confer resistance either directly (*97*) or additively with other variants (*21*) in *C. elegans*. Neither gene contained high effect variants, however, both contained a number of non-synonymous variants, including three moderately differentiated variants between pre- and post-treatment in *osm-1* (pos. 3,109,316 [Gln1540Pro], $P$ = 8.444e-05; pos. 3,109,310 [Val1542Gly], $P$ = 8.337e-06; pos. 3,109,002 [Glu1577Asp], $P$ = 4.936e-04) and four variants with significant but less differentiated variants in *che-11*. A glutamate transporter family protein *glt-5* (HCON_00035710) was also identified; however, there was no evidence of selection at this locus.

<u>Half-dose followed by double-dose ivermectin treatment primes selection at novel loci</u>

The X-QTL experiments of ivermectin selection used a standard treatment dosage of 0.2 mg/kg of ivermectin per host weight. In an attempt to improve the resolution of the QTL identified, and perhaps, refine the variants under selection to those that specifically confer high levels of resistance, we performed a follow-up experiment to the X-QTL using twice the standard dose treatment (0.4 mg/kg). To do so, we first treated sheep infected with the F3 generation of the cross with a half-standard dose (0.1 mg/kg), followed by a subsequent double-standard dose treatment (0.4 mg/kg) 14 days later. Similar to the X-QTL experiments, larvae were sampled pre- and post- each treatment as before (pools of 200 $L_3$), and included a time-matched, untreated control.

Relative to timed matched controls (**fig. S10 A and B**), at the half standard dose we identified discrete regions of chromosomes 2 and 5 (**fig. S10 C)** that increased in genetic differentiation after the double dose treatment (**fig. S10 D**). After replication, however, different patterns of selection emerged; although the chromosome 5 region of differentiation was consistent between all replicates, we identify replicate-specific QTL in the population treated with double the standard dose of ivermectin on chromosomes 1, 2, 3, and 4 (**fig. S10 E, F,** and **G** show biological replicates, with new QTL highlighted by a red, dashed box). There is also variation that may suggest selection at additional loci, however, they cannot be statistically differentiated from the high degree of stochastic background genetic variation throughout the genome in these pooled samples. One hypothesis is that these QTLs represent stochastic noise unrelated to drug exposure, evidenced by the fact that they only appear in a single replicate (apart from the chromosome 2 peak that does replicate from the X-QTL analyses). Alternatively, these discrete QTL may provide evidence of selection on different genes in different sub-populations that contribute to the resistance phenotype (i.e. additively to provide additional levels of resistance and/or to compensate for fitness costs associated with variation needed for resistance) together with the necessary and sufficient causative allele on chromosome 5. If the latter is correct, these data allow us to speculate that underdosing of drug treatment to control parasites like *H. contortus* (represented here by the half-standard dose treatment) may "prime" different loci providing low-level but nonetheless improved fitness in the presence of the drug, and may, in turn, lead to different genome-wide signatures of selection in field populations that have been exposed to different management strategies used to control them.

The relative contribution of these secondary QTL towards the generation of a resistance phenotype, either by their effect on relative fitness (i.e. selection coefficient) or whether they occur independently of one another, is difficult to determine precisely using the Pool-seq approach applied here. It may be possible to resolve these further by refining the genetic crossing strategy used here by, for example, the use of larger sampled population sizes or later generations to increase the recombinant fraction and thereby decrease the amount of linked variation co-inherited with causal variants. Alternative approaches could also include decreasing the number of individuals used to initiate the genetic crosses (for example, using single resistant females with multiple males (52)) with multiple replicates of each cross to increase the likelihood of initiating a genetic cross with one or another secondary QTL but not multiple QTLs. We acknowledge that these alternative approaches are coupled with additional technical challenges and associated increases in cost, including but not limited to: a lower efficiency / higher failure rate of single worm infections (relative to the bulk worm infections we describe); the need for additional host animals to establish such infections - the lower efficiency of initiating an infection may, therefore, require multiple independent infections, and at least one infection per QTL would be required (as opposed to the triplicate sheep for all QTLs we describe); and the challenge of maintaining secondary QTLs, which may become lost via genetic drift over multiple generations, without understanding the precise selection pressure by which they are maintained.

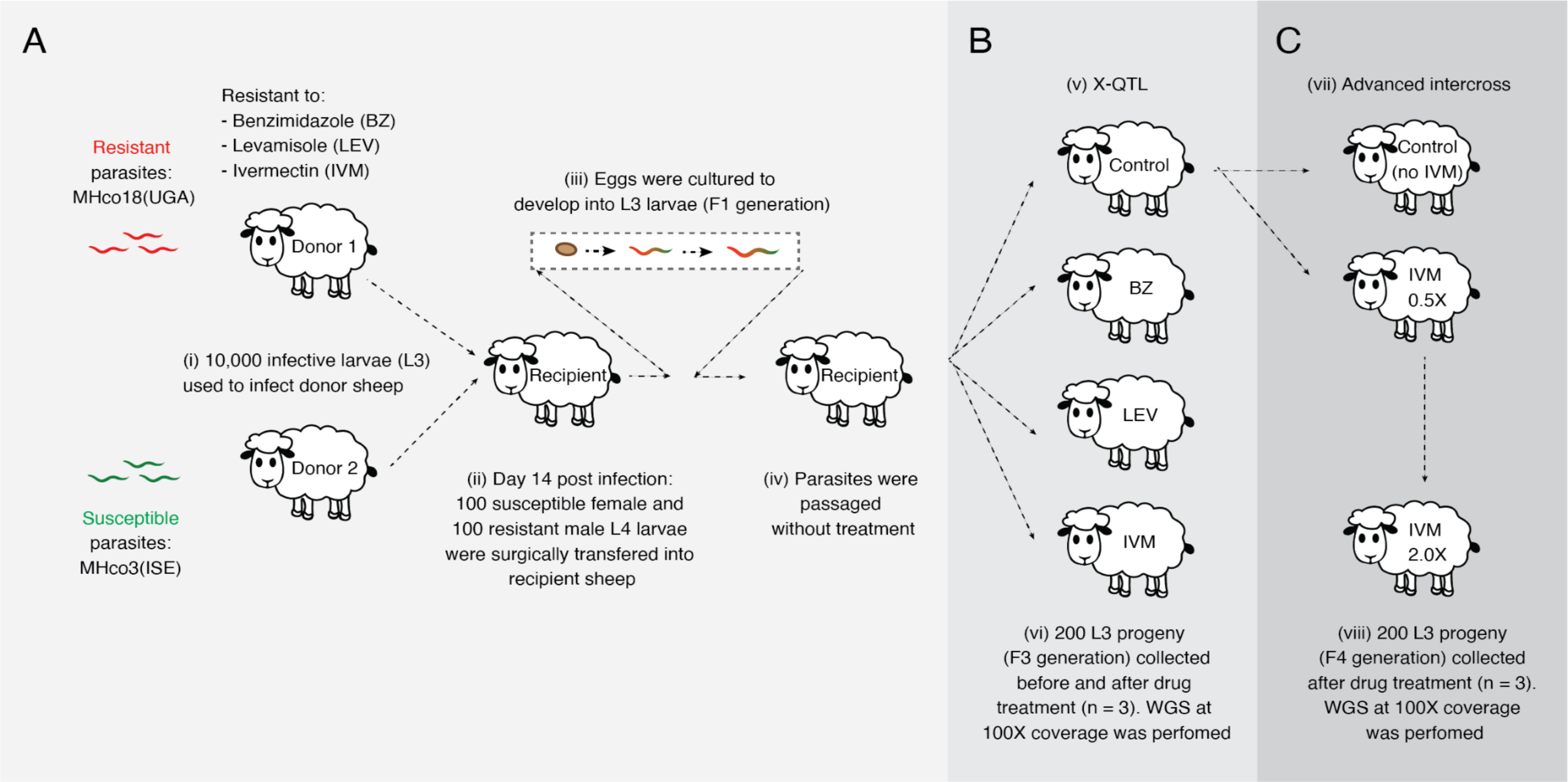


**Fig. S1. Outline of the genetic cross, X-QTL, and advanced intercross sampling design.** (**A**) A genetic cross between the anthelmintic susceptible MHco3(ISE) and multi-drug resistant MHco18(UGA) was used to map genetic loci associated with resistance to benzimidazole, levamisole and ivermectin drug treatment. (**B**) A drug selection X-QTL experiment was performed on the F2 generation, which was split into four groups that were exposed to either (i) benzimidazole, (ii) levamisole, (iii) ivermectin, or (iv) remained untreated. (**C**) An advanced intercross experiment using F3 generation progeny pooled from pre-treated (untreated) $L_3$ were subjected to half- followed by double-standard doses of ivermectin. For both the drug selection X-QTL and advanced intercross experiments, pools of $L_3$ (n = 200) were collected both pre- and post-treatment from drug-exposed and time-matched untreated controls. Each experiment was performed in triplicate. Whole-genome sequencing was performed on the pools, after which genetic diversity between pre- and post-treatment was compared.

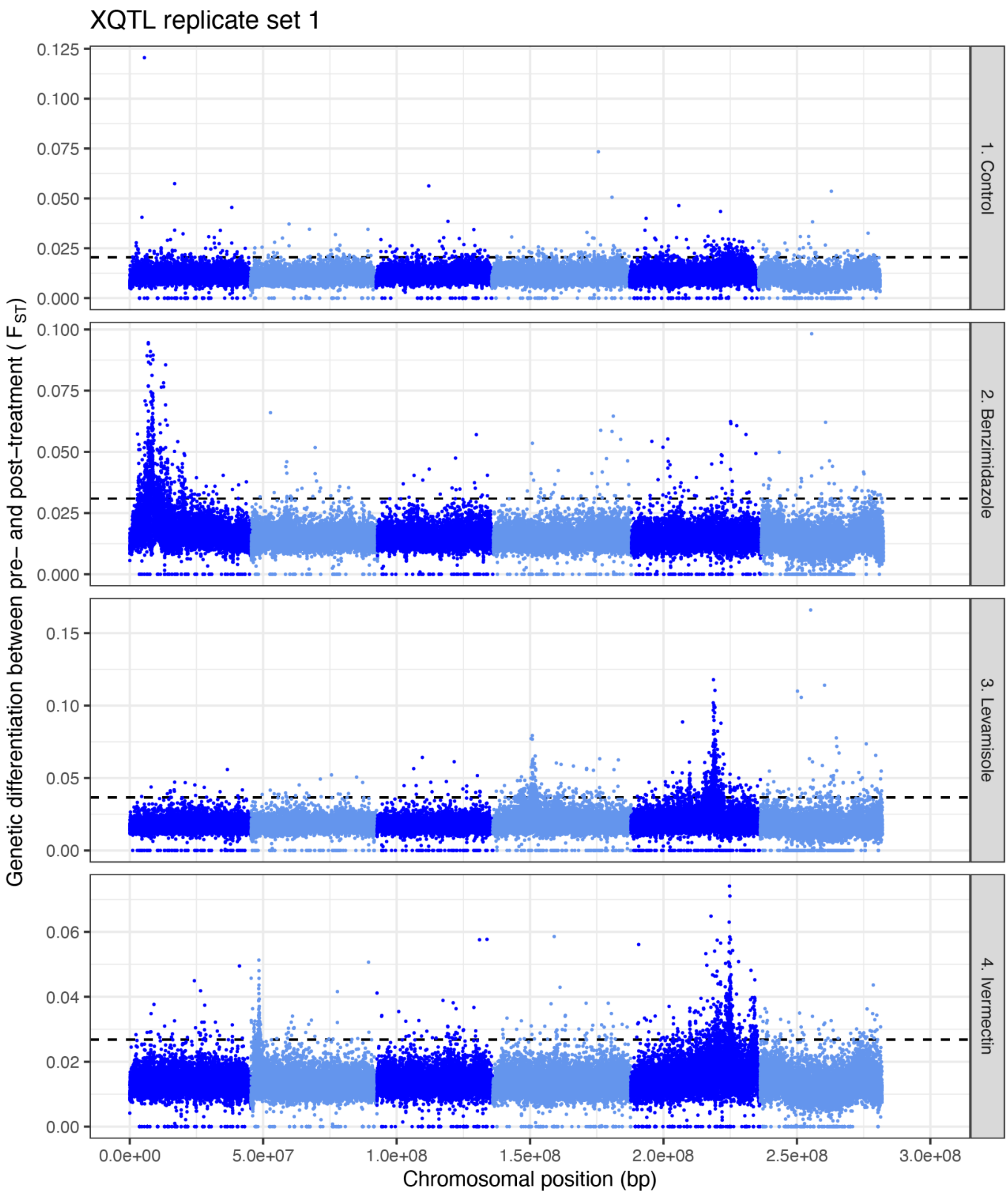

XQTL replicate set 1
Genetic differentiation between pre- and post-treatment ( $F_{ST}$)
1. Control
2. Benzimidazole
3. Levamisole
4. Ivermectin
Chromosomal position (bp)
0.0e+00
5.0e+07
1.0e+08
1.5e+08
2.0e+08
2.5e+08
3.0e+08

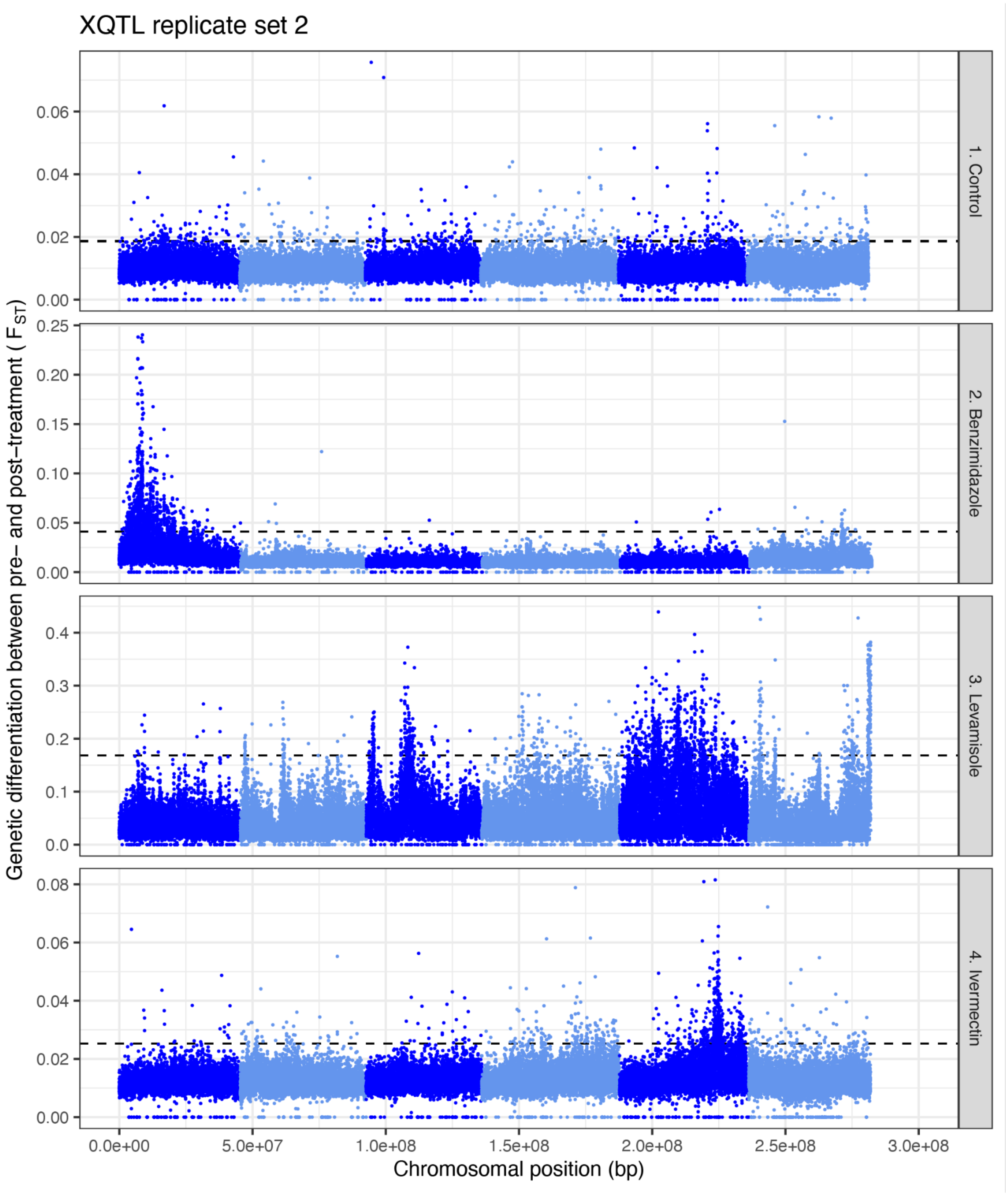

XQTL replicate set 2
1. Control
2. Benzimidazole
3. Levamisole
4. Ivermectin
Genetic differentiation between pre- and post-treatment ( $F_{ST}$)
Chromosomal position (bp)
0.0e+00
5.0e+07
1.0e+08
1.5e+08
2.0e+08
2.5e+08
3.0e+08

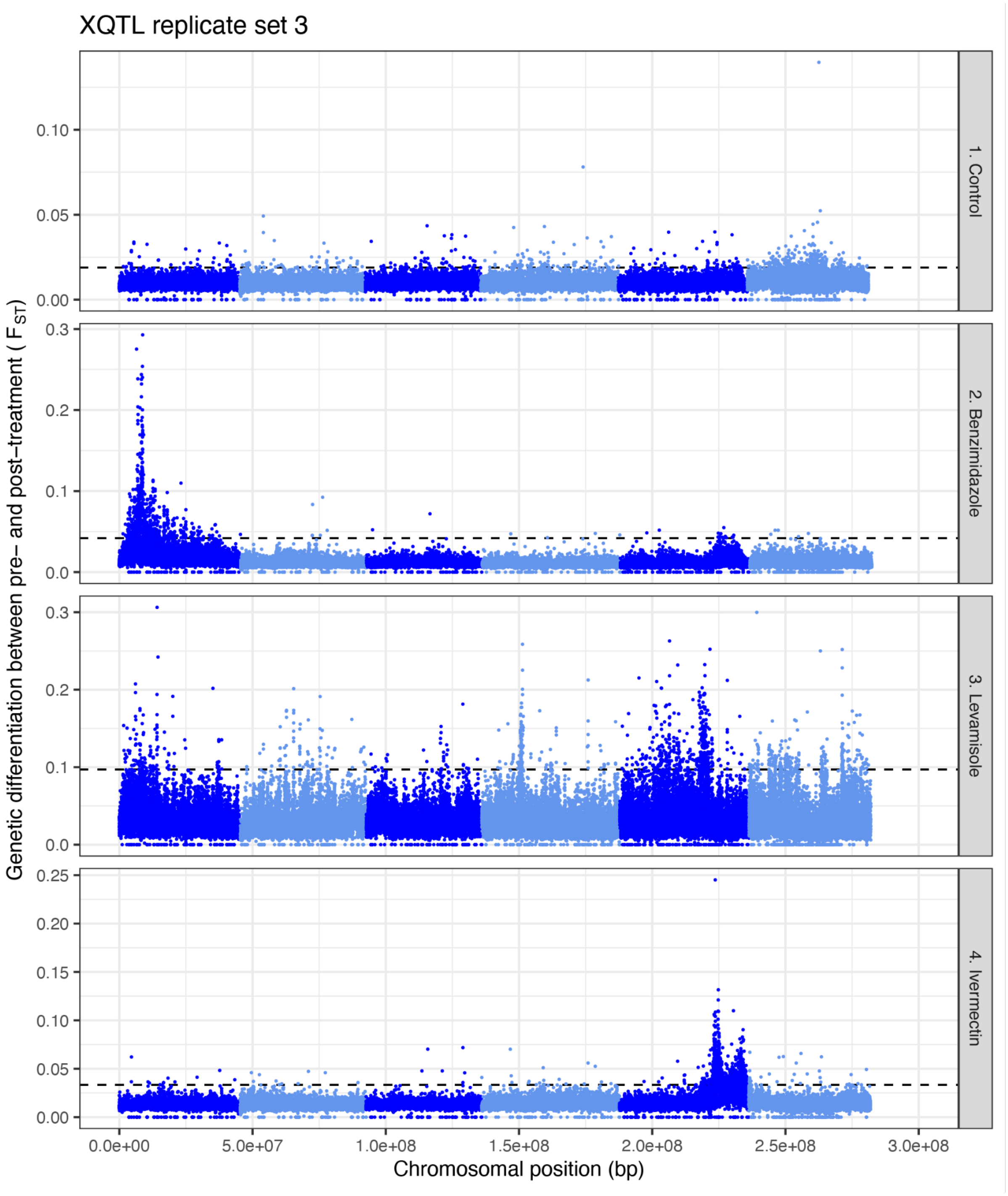


**Fig. S2. Genome-wide differentiation ($F_{ST}$) between pre- and post-treatment populations of three independent replicate sets of the X-QTL experiment.**

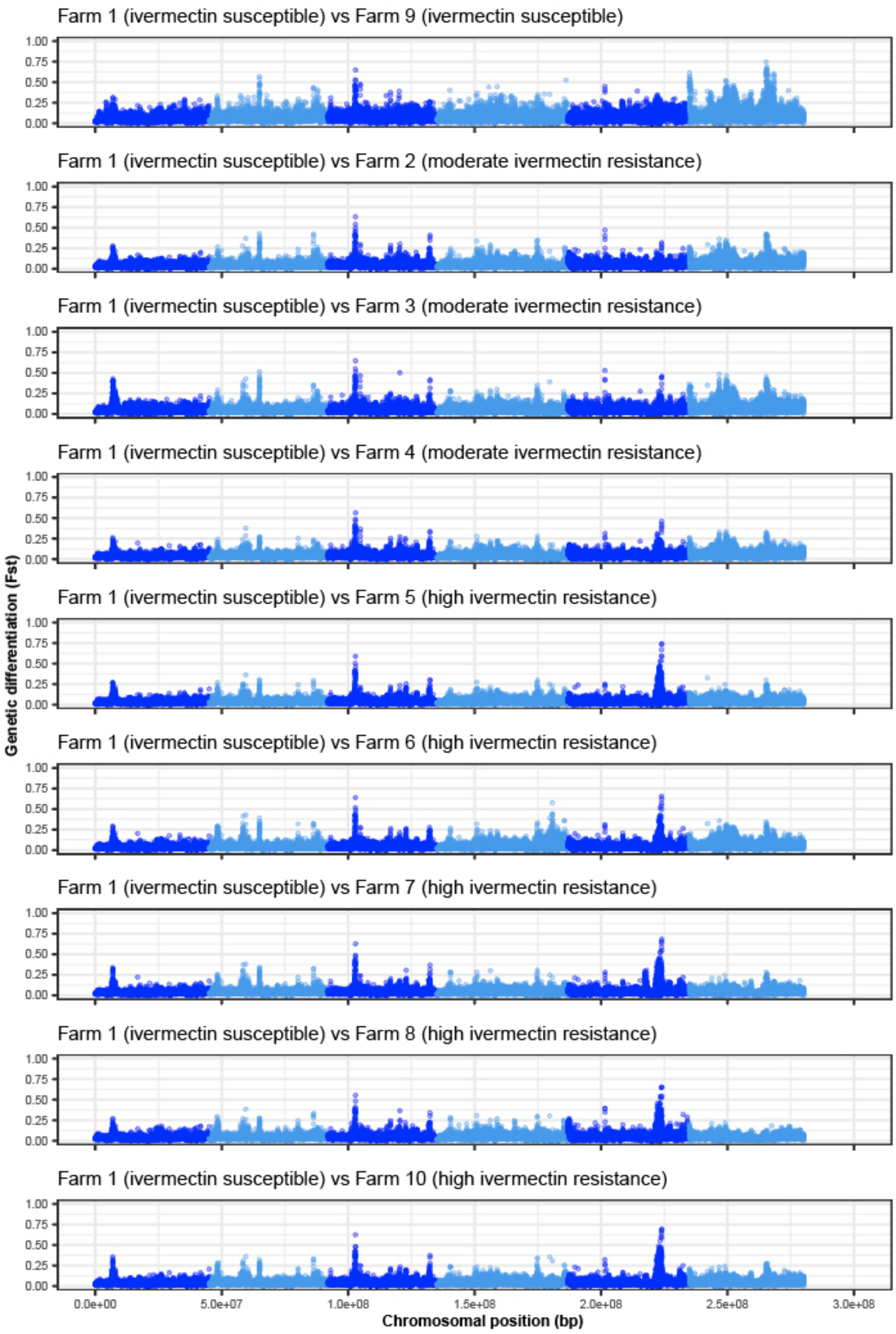


**Fig. S3. Pairwise comparison of differentiation ($F_{ST}$) between a susceptible laboratory *H. contortus* isolate (Farm 1) and US farms that differ in sensitivity to ivermectin.** Farm 9 is susceptible and the rest have been classified as being moderately or highly ivermectin resistant, based on the DrenchRite larval development assay.

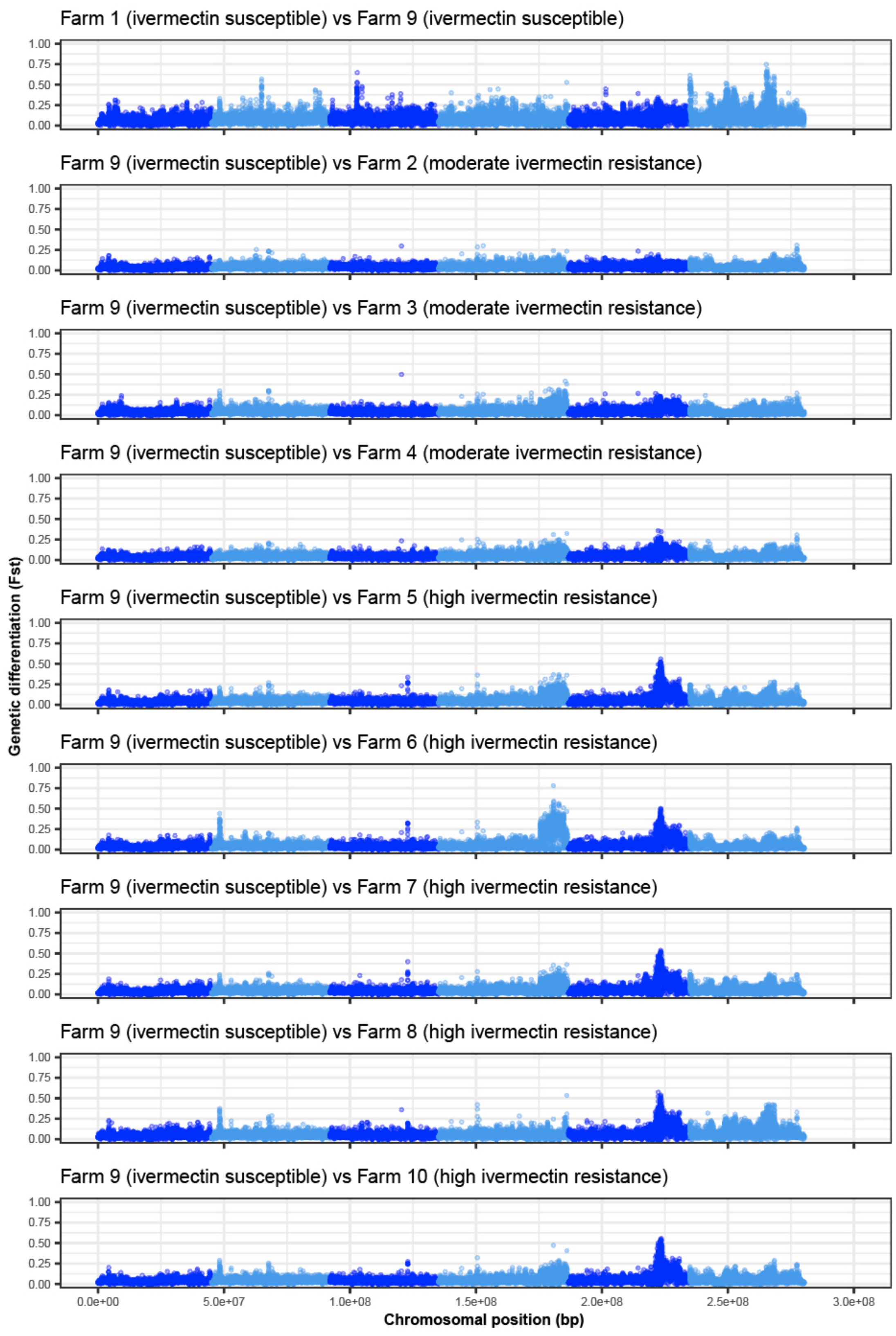


**Fig. S4. Pairwise comparison of differentiation ($F_{ST}$) between a susceptible laboratory *H. contortus* isolate (Farm 9) and US farms that differ in sensitivity to ivermectin.**

A

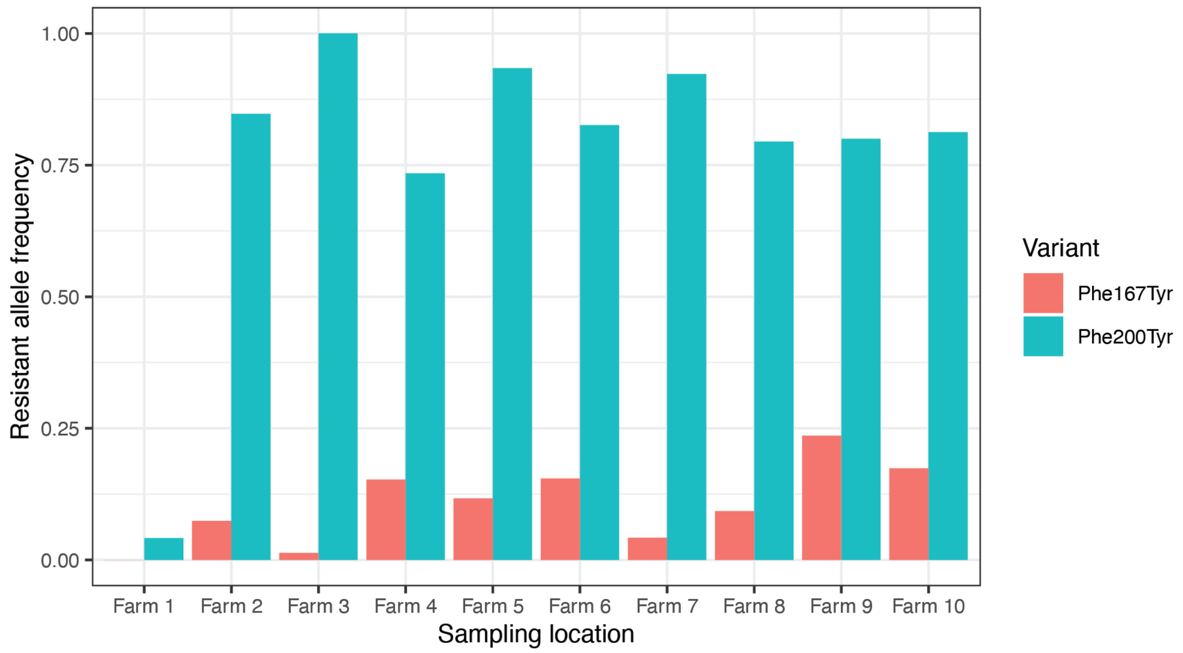


**Fig. S5. Frequency of Phe167Tyr and Phe200Tyr variants of beta-tubulin isotype 1 associated with benzimidazole resistance from a US laboratory strain and 9 US farm isolates.** The frequency of Phe167Tyr and Phe200Tyr variants were estimated using whole genome sequencing data from *H. contortus* $L_3$ on 10 US farms. DrenchRite larval development assays demonstrate that worms from Farm 1 are susceptible to benzimidazole treatment, consistent with the absence of resistance-associated variants. However, all other farms are phenotypically resistant to benzimidazole anthelmintics (**Table S2**).

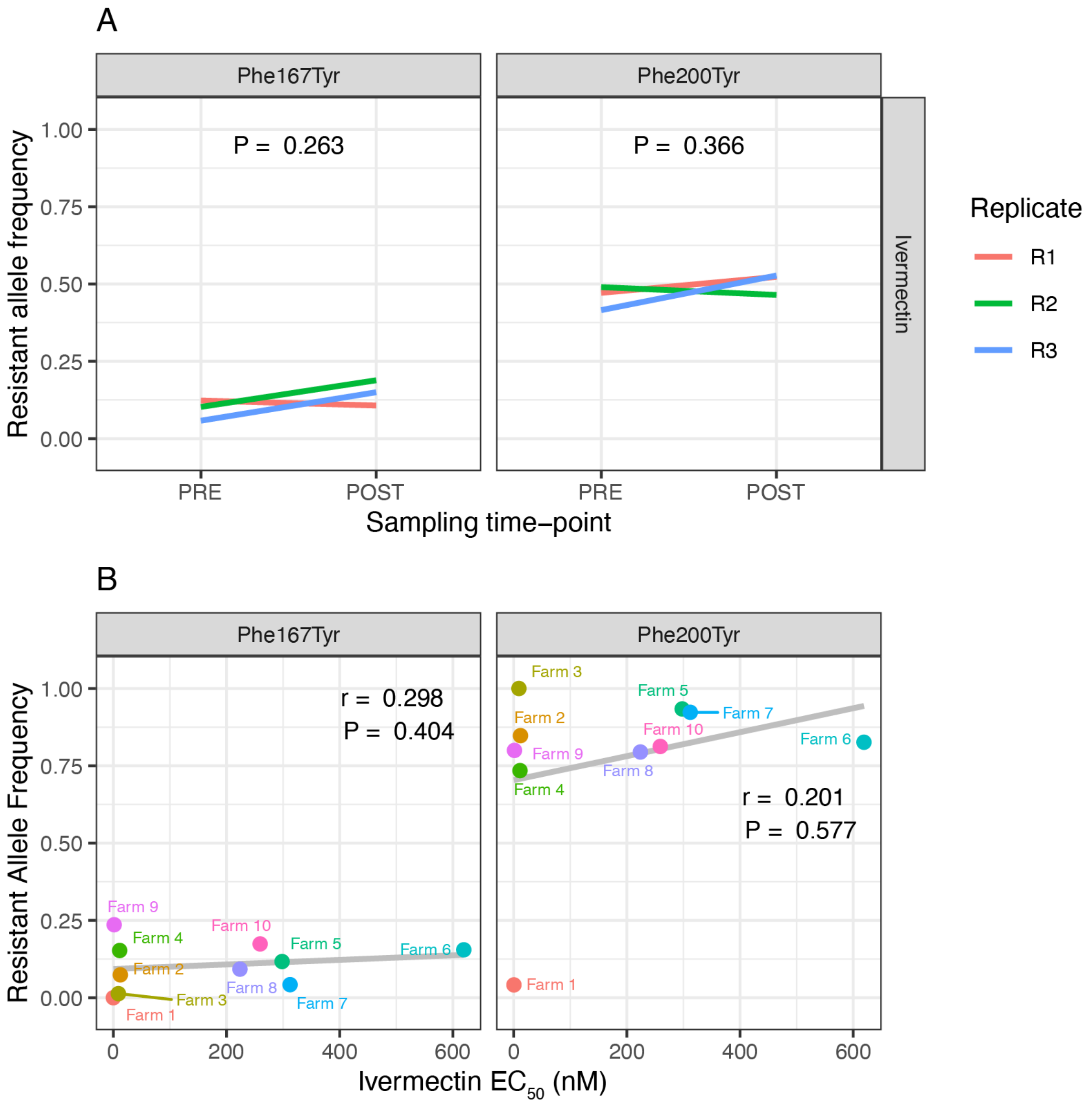


**Fig. S6. No evidence of ivermectin selection on beta-tubulin isotype 1 resistant alleles in the genetic cross or US farms.** (**A**) Estimation of Phe167Tyr and Phe200Tyr variant frequencies of *H. contortus* $L_3$ collected from pre- and post-ivermectin treated animals of the cross. Neither variant showed a significant change in variant frequency, measured using a paired t-test from which the *P*-values are shown. (**B**) Estimation of Phe167Tyr and Phe200Tyr variant frequencies from *H. contortus* $L_3$ and their correlation with DrenchRite measurement of ivermectin resistance on 10 US farms. For both variants, no significant correlations were identified. Pearson's correlation (r) and associated *P*-value together with the trendline of the linear regression are shown.

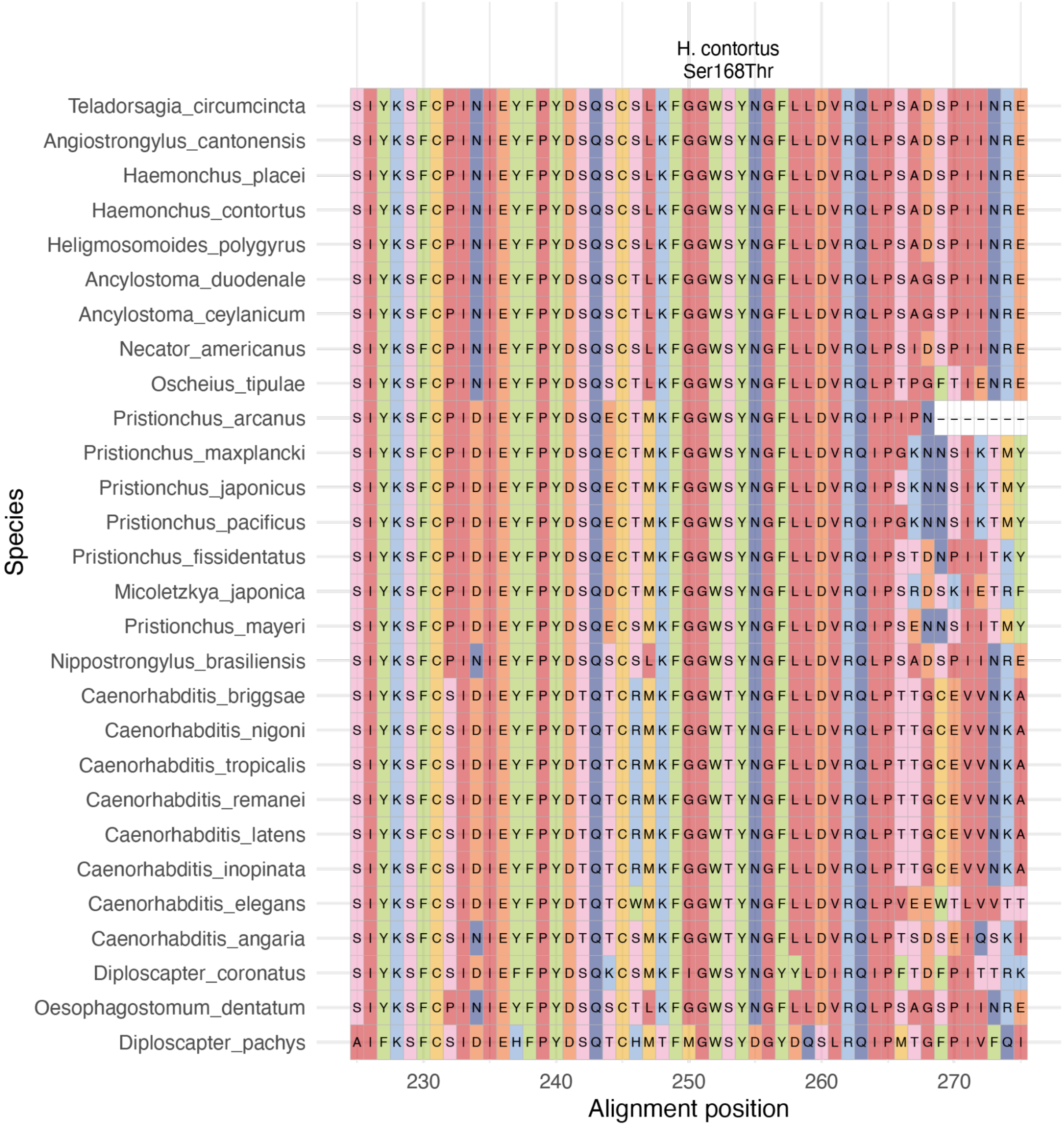


**Fig. S7. Multiple sequence alignment of Clade V orthologs of acetylcholine receptor subunit ACR-8.** A Ser167Thr variant in *Haemonchus contortus* (Ser position highlighted) is correlated with levamisole resistance. This multiple sequence alignment focused on alignment positions 225-275 surrounding the *H. contortus* Ser167Thr position demonstrates high sequence conservation of the Ser residue among all Clade V nematodes analysed, except for species within the *Caenorhabditis* genus (which contain Thr).

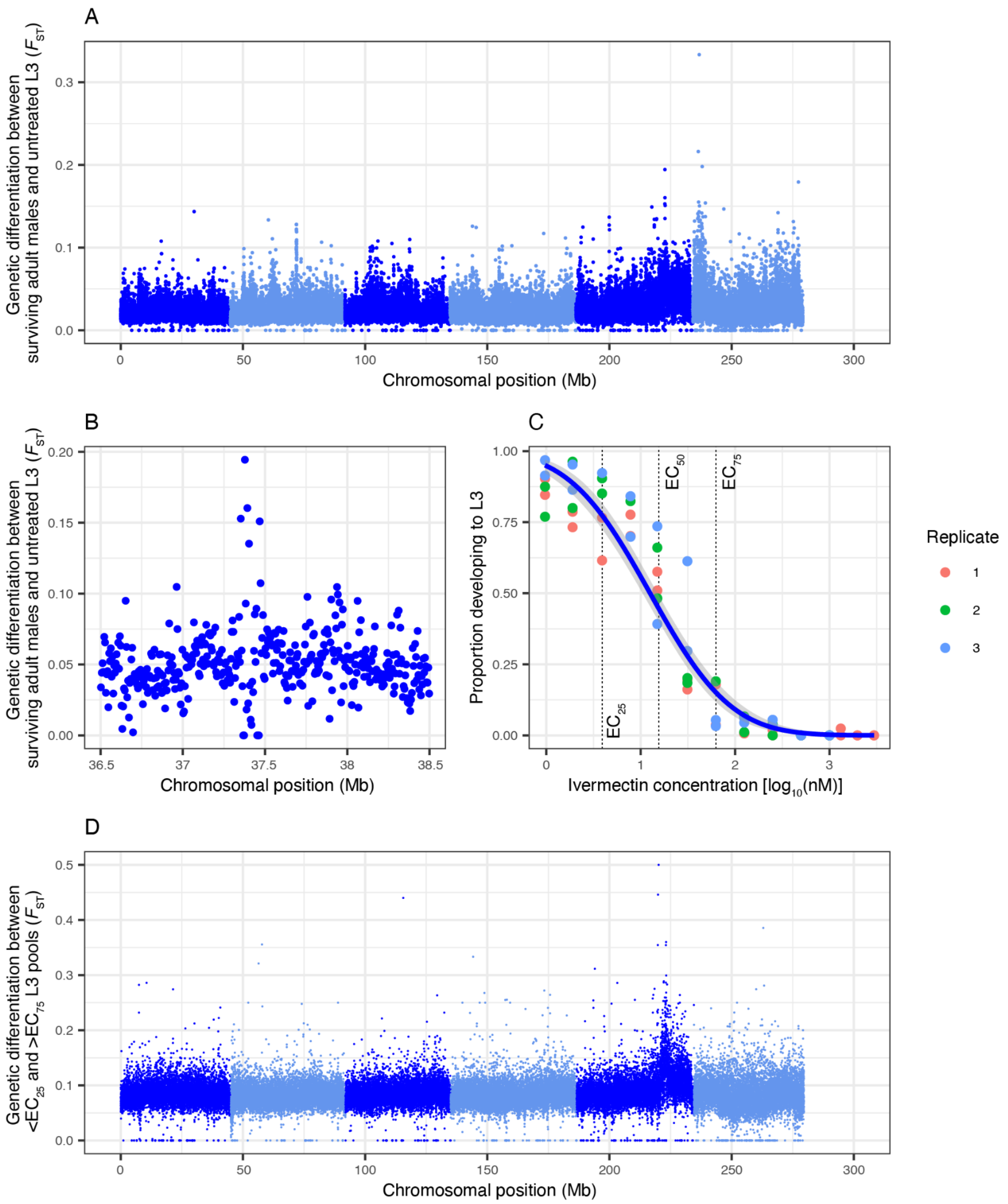


**Fig. S8. Consistent evidence of selection on the chromosome 5 locus with respect to direct versus indirect measurements of genetic diversity after treatment with ivermectin.** (**A**) Genome-wide analysis of genetic differentiation between pre-treatment L3 (n = 200; recovered from host faeces) and adult males (n = 40; recovered by host necropsy) directly exposed to and having survived treatment with ivermectin. (**B**) The same genomic interval on chromosome 5, as shown in Figure 4 A, highlights a smaller peak region. (**C**) Dose-response curve of ivermectin using larval development assays to determine the $EC_{50}$. (**D**) Genome-wide analysis of genetic differentiation between poorly developing $L_1/L_2$ stage larvae at $EC_{25}$ and developing $L_3$ at $EC_{75}$.

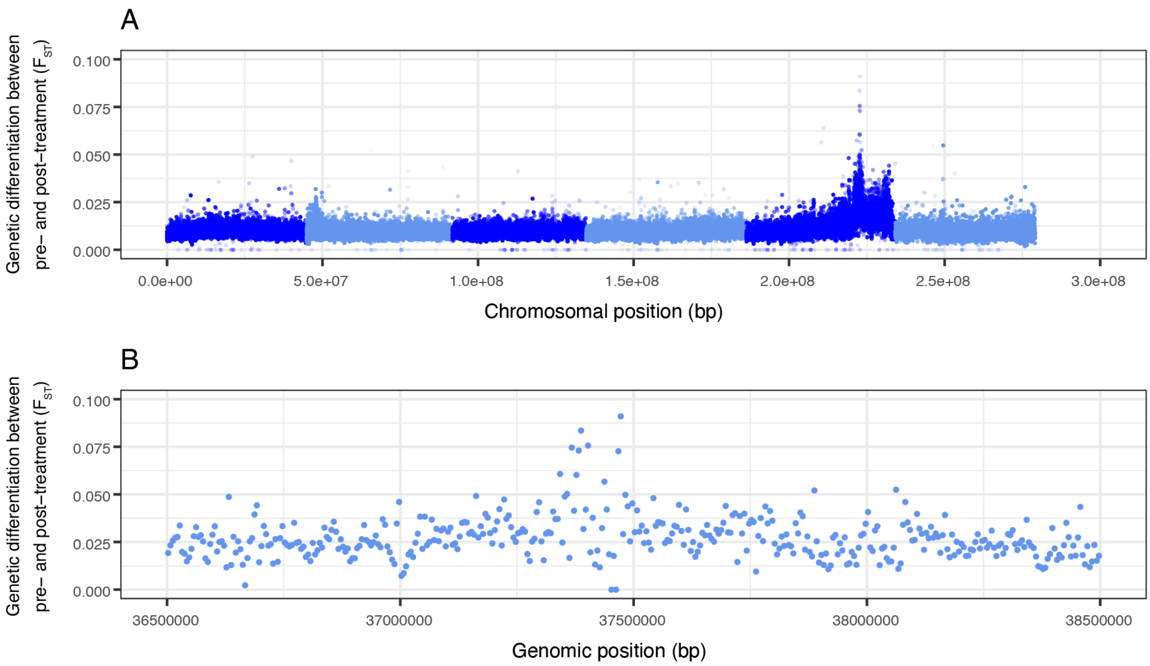


**Fig. S9. A higher number of $L_3$ per pool helps to increase the resolution of the peaks of differentiation.** (**A**) Whole-genome comparison of pools of $L_3$ (n = ~5000) pre- and post-ivermectin treatment. (**B**) The same genomic interval on chromosome 5 as shown in Figure 5 B (i.e., 36.5 to 38.6 Mb), highlighting the greater resolution of the ivermectin QTL as a consequence of more recombinant individuals in the $L_3$ pooled (n = ~5000) relative to the X-QTL analysis (n = 200 $L_3$ per pool).

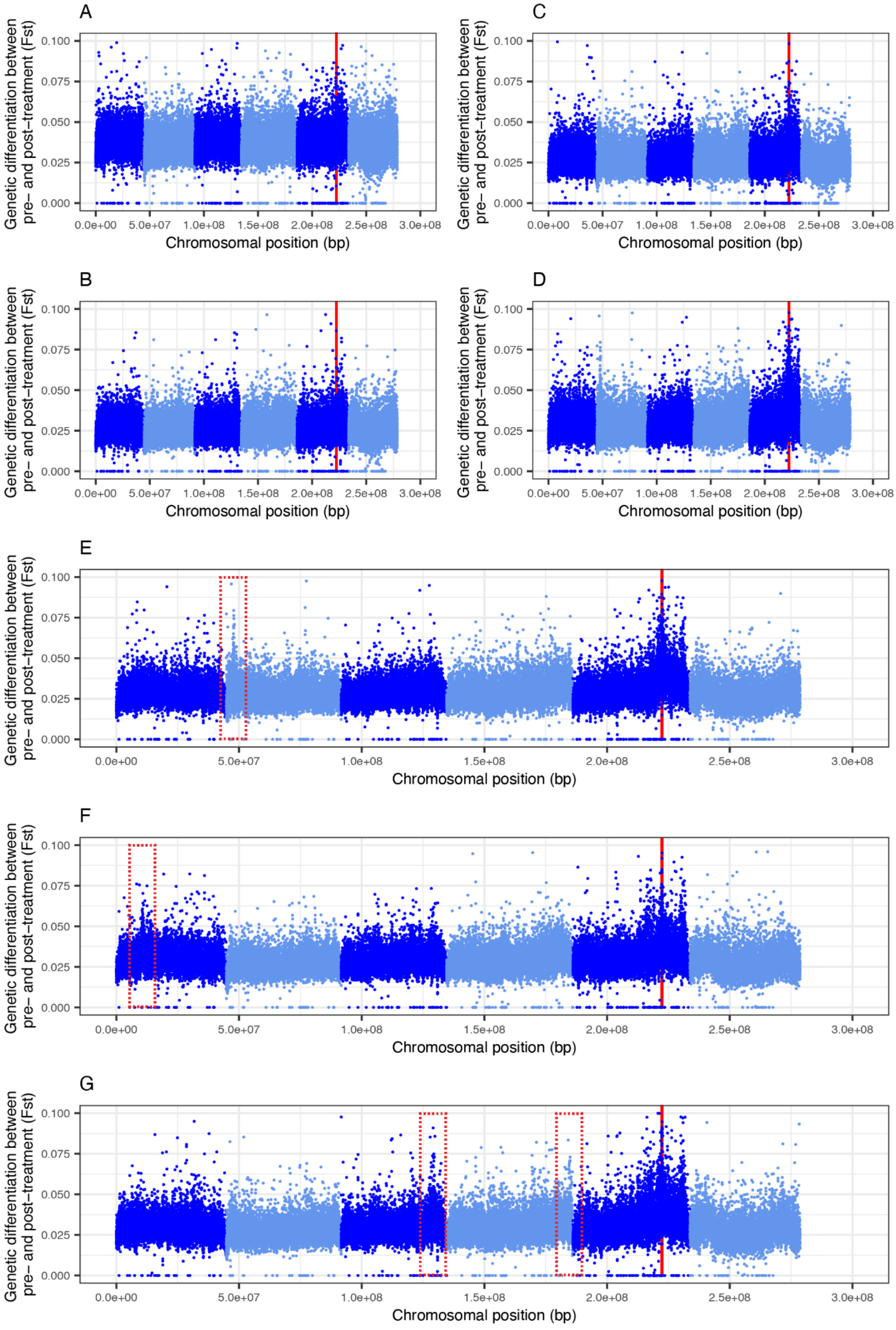
A
B
C
D
E
F
G
Genetic differentiation between pre- and post-treatment (Fst)
0.000
0.025
0.050
0.075
0.100
0.0e+00
5.0e+07
1.0e+08
1.5e+08
2.0e+08
2.5e+08
3.0e+08
Chromosomal position (bp)

**Fig. S10. Differential dosing changes the genetic signature of ivermectin treatment.** Comparison of untreated (**A**), time-matched control (**B**) and ivermectin treated populations that were sampled before and after the treated population were exposed to half-dose (**C**) followed by double dose (**D**) standard ivermectin treatment. When the experiment is replicated (each plot **E, F,** and **G** shows an individual replicate), new discrete QTL are identified (red dashed boxes) in the double-dose treatment comparisons, in addition to the primary QTL for ivermectin resistance on chromosome 5 (solid red line) that appears in all experiments in which ivermectin selection is applied.
